# Augmenting the Bayesian Brain with learned and reusable world-model components for flexible cognition

**DOI:** 10.64898/2026.05.06.722922

**Authors:** Charles Findling, Junseok K. Lee, Jacob J. W. Bakermans, Alexandre Pouget, Valentin Wyart

**Affiliations:** Department of Basic Neurosciences, University of Geneva, Switzerland; Laboratoire de Neurosciences Cognitives et Computationnelles (LNC2), Institut National de la Santé et de la Recherche Médicale (Inserm), Paris, France; Département d’Études Cognitives, École Normale Supérieure – Université PSL, Paris, France; Sainsbury Wellcome Centre for Neural Circuits and Behaviour, University College London, London, UK

## Abstract

The Bayesian Brain hypothesis assumes that cognition relies on internal generative models of the world, yet existing implementations remain constrained by pre-specified, task-specific generative structures and computationally heavy iterative inference schemes. Here, we introduce modular neural state-space models as a scalable realization of the Bayesian Brain, replacing fixed generative structures and pre-specified inference rules with learned world-model components and amortized neural updates. This framework preserves the core commitment to explaining observations through hidden causes while making inference learned and reusable rather than pre-specified and task-specific. Our modular implementation of these models affords learned components to be seamlessly recombined and stacked across superficially different tasks that share similar latent dynamics. Such computational reuse supports zero-shot generalization and predicts selective correlations of inference parameters between tasks. We confirm these key predictions in human behavior, identifying learned and reusable world-model components as a candidate computational principle for flexible cognition.

## Introduction

A central idea in cognitive neuroscience is that the brain maintains internal generative models to interpret and predict its sensory input^1–5^. This **Bayesian Brain** framework has been influential across perception, learning, reasoning and decision-making^6–9^. Yet, and despite its conceptual appeal, algorithmic realizations of this framework have remained constrained by two longstanding limitations. First, they typically rely on fixed, pre-specified generative structures^10–12^, offering little capacity for meta-learning, for discovering new latent variables, or for reusing learned structure across tasks. As a result, these models provide only a limited account of how knowledge acquired in one setting may transfer to novel situations that share the same latent dynamics. Second, inference rapidly becomes intractable in richer conditions, forcing models to assume overly simple, low-dimensional dynamics or to rely on complex variational or sampling procedures. These inference schemes are inherently iterative – requiring repeated updates, sampling, or optimization^13–15^ – raising both computational and biological implementation challenges. Together, these constraints have contributed to the view that the Bayesian Brain framework applies only in a restricted set of real-world conditions.

Here, we overcome these limitations by augmenting the Bayesian Brain with **neural state-space models (SSMs)** – models that represent hidden states of the environment while using neural networks to learn components of the generative process from experience. Neural SSMs replace both fixed, pre-specified generative structures and inference rules. Instead of assuming a particular generative model, they acquire flexible world-model components directly from experience. And instead of requiring iterative procedures such as variational methods, expectation-maximization or sampling, they implement amortized inference by learning to update their beliefs directly through neural dynamics^16,17^. This provides a simple neural mechanism for belief updating and a biologically plausible alternative to explicit inference schemes. Crucially, neural SSMs are trained to fit a generative model of the world to observations, preserving the core Bayesian aim of explaining sensory input through hidden causes while avoiding the computational bottlenecks of explicit inference. The resulting framework is both tractable and expressive, enabling probabilistic computation in real-world conditions that challenge existing approaches.

A second central contribution is that neural SSMs can be structured in a modular way. As we will show, by assigning different aspects of the generative process (world model) to distinct neural components, the architecture introduces strong inductive biases toward a modular world-model structure. These components define a repertoire of cognitive strategies that can be re-used across conditions that share latent generative properties, even when the statistics of observations (but also the nature of observations and of associated latent states) differ^18^. This modularity also allows previously learned neural components to be stacked and recombined to solve more complex tasks that were never encountered during training. In this way, neural SSMs support zero-shot composition of learned world-model components, rather than requiring each new task to be learned from scratch. This stands in contrast to large end-to-end networks trained purely for predictive accuracy or performance. Indeed, such models lack explicit generative factorization and therefore show limited computational ‘recycling’ and minimal zero-shot transfer across tasks^19–22^.

If the human brain implements such modular neural SSMs, then the reuse of world-model components should leave measurable and selective signatures in behavior. In particular, individuals should display correlated learning patterns across tasks that are superficially dissimilar but share the same latent generative structure. By contrast, such correlations should be absent or markedly reduced between tasks that do not recruit the same latent structure, even when they are superficially similar. Importantly, the reuse of a world-model component does not imply identical behavior across tasks, but it predicts significant cross-task relationships in specific aspects of behavior that depend on the world-model component being reused across tasks.

In this work, we use neural SSMs to formalize this modular world-model framework, demonstrate its capacity for zero-shot recombination of learned components, and test the behavioral predictions of computational reuse in human behavior. Together, our results support the view that the brain combines generative world modeling with reusable amortized computations to achieve structured inference and cognitive flexibility.

## Results

### Neural state-space models learn generative world models in cognitive tasks

As a first step, we asked whether neural SSMs can serve as a computational realization of the Bayesian Brain – that is, whether they can learn generative models of the world that explain observations through hidden causes – beyond the regime where classical implementations (e.g., linear Gaussian or discrete-state models with known parameters) remain tractable. To this end, we examined two canonical forms of learning that are central to cognitive neuroscience yet already challenge classical Bayesian approaches: **reversal learning**, which requires tracking changes in latent contingencies, and **associative learning**, which requires discovering and maintaining structured relationships between cues and stimuli.

We first considered a **two-armed bandit task** that instantiates **reversal learning** under two simultaneous sources of uncertainty (**Fig. 1a**; see **Methods**). On each trial, the agent chooses between two symbols (arms), and receives a continuously-valued reward (outcome) sampled from a probability distribution associated with the chosen symbol. The goal is to sample from the better arm – associated with the higher mean reward – whose identity can change through reversals controlled by a transition structure (**Fig. 1a**). An example instance of the two-armed bandit task is depicted in **Fig. 1b**, illustrating reversals in the latent state (i.e., the identity of the better arm) and the rewards received by an agent. Although the task appears simple, the agent must: 1. monitor the identity of the better arm (**latent state**), 2. estimate a latent state volatility governing the rate of reversals (**transition uncertainty**), and 3. estimate the sampling distribution of rewards (**emission uncertainty**). Together, these interacting sources of uncertainty define a hierarchical generative problem for which exact Bayesian updates are unavailable and standard solutions rely on iterative, task-specific inference schemes.

**Figure 1.**
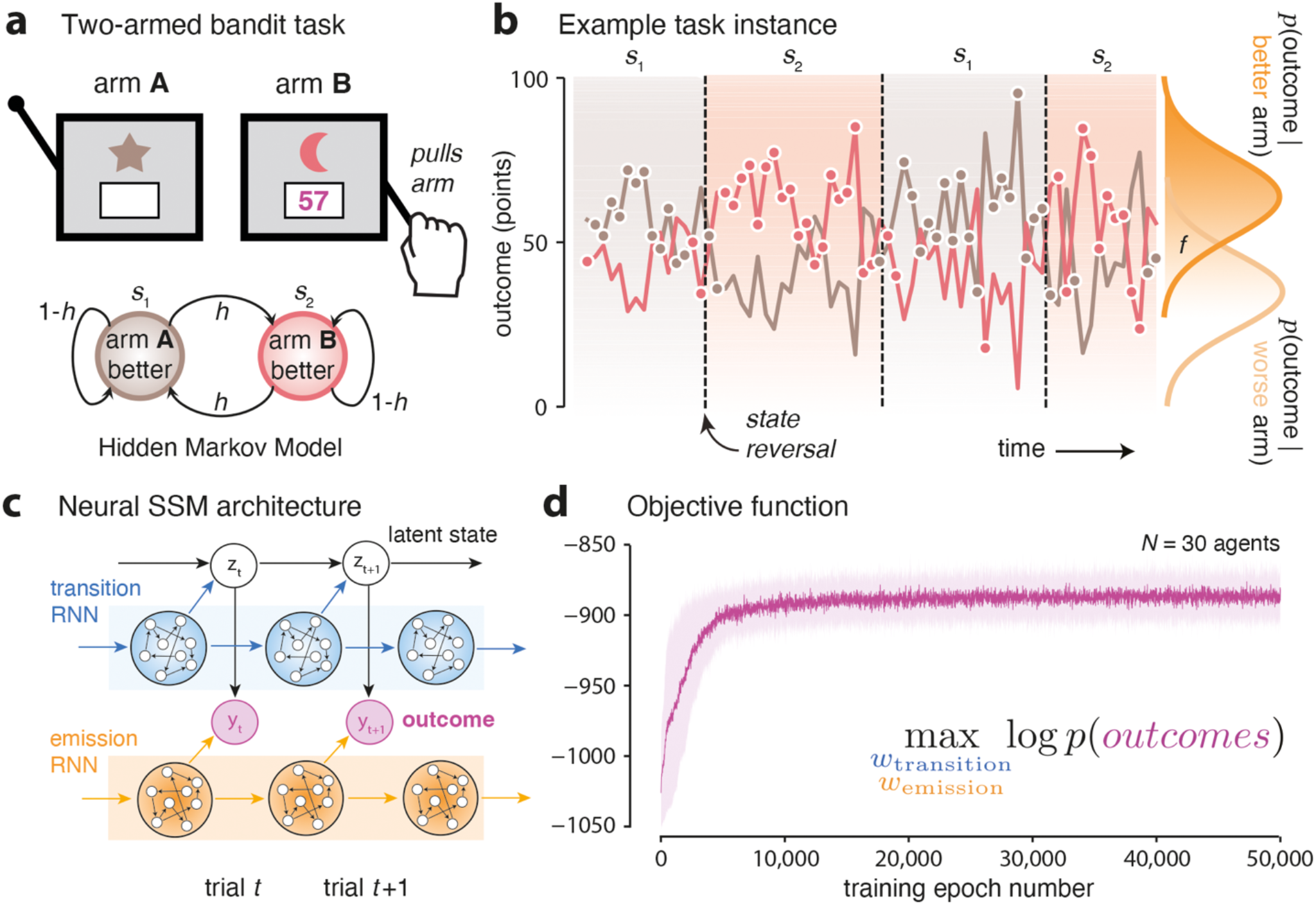
Neural state-space models applied to reversal learning. (**a**) Two-armed bandit task with reversals. On each trial, the agent selects between arm A and arm B and receives a continuous-valued outcome. A latent binary state determines which arm is currently better. The latent state evolves according to a Markov process with reversal probability *h*, governing transition uncertainty (state volatility). Outcomes are generated from noisy reward distributions conditioned on whether the chosen option is currently the better or worse one, introducing emission uncertainty in the mapping between latent reward state and observed feedback. (**b**) Example task instance. Trial-by-trial reward trajectories for both arms are shown across time. The latent state (identity of the better arm) undergoes unsignaled reversals (dashed vertical lines). Outcomes for both arms are displayed for illustration, although the agent observes only the outcome corresponding to its chosen arm. Circle markers indicate the actions selected by an example neural SSM agent, which chooses the arm with the highest posterior probability of being most-rewarding at each trial. (**c**) Neural SSM architecture for reversal learning. The model comprises a discrete latent state *z*_*t*_ encoding the currently higher-rewarding arm and two recurrent components that estimate emission and transition parameters from experience. Inference over the latent state is performed exactly via likelihood recursion. The emission module outputs the probability distribution *p*(*y*_*t*_| *a*_*t*_= *z*_*t*_), with *y*_*t*_the outcome and *a*_*t*_ the chosen action. The transition component outputs the state volatility *h*_*t*_. Thus, emission and transition uncertainties are updated through amortized recurrent dynamics, while belief updating over the latent state remains analytically exact. (**d**) Training dynamics. Learning curve showing the marginal log-likelihood of observed outcomes as a function of training epochs (*N* = 30 independently trained neural SSMs). Parameters of both transition and emission networks are optimized jointly to maximize *log p*(*y*_1:*T*_), demonstrating stable convergence.

We capture this structure using neural SSMs that jointly learn a **transition network** and an **emission network** from experience, implementing inference over latent parameters through amortized neural updates rather than explicit iterative procedures (**Fig. 1c**; see **Methods**). Inference over the latent state *z*_*t*_itself is performed exactly using likelihood recursion^23^ (see **Methods**), yielding analytically tractable belief updates. On each trial, the emission network updates its internal state based on the current posterior belief over the selected arm and the obtained reward, allowing it to learn the sampling distribution of rewards from experience. In parallel, the transition network receives an evidence signal derived from the emission network – specifically, the log-likelihood ratio between the hypothesis that the chosen arm is the better one and the alternative hypothesis – and uses it to update its estimate of state volatility. Unlike standard training approaches optimizing task performance, training of neural SSMs is performed by maximizing the likelihood of observed outcomes (rewards in this case), as predicted by the Bayesian Brain framework, and converges reliably across *N* = 30 neural SSMs trained independently (**Fig. 1d**).

After training, neural SSMs reproduce key behavioral signatures of adaptive learning under uncertainty (**Fig. 2a**). Performance decreases as environmental volatility increases (**Fig. 2a**, top), reflecting the growing difficulty of tracking rapidly changing reward contingencies. Performance likewise declines as outcome reliability decreases (**Fig. 2a**, bottom), where outcome reliability is quantified by the false-feedback rate *f* – the amount of overlap between the sampling distributions of rewards associated with the better and worse arms. These performance profiles closely match those of the optimal Bayesian agent (**Fig. 2a**). Across task conditions, choice accuracy reaches 81.1% for neural SSMs, compared to 81.7% for the optimal Bayesian model – which relies on iterative and computationally demanding inference, implemented here using particle filtering (see **Methods**; **Supplementary Table 1**).

**Figure 2.**
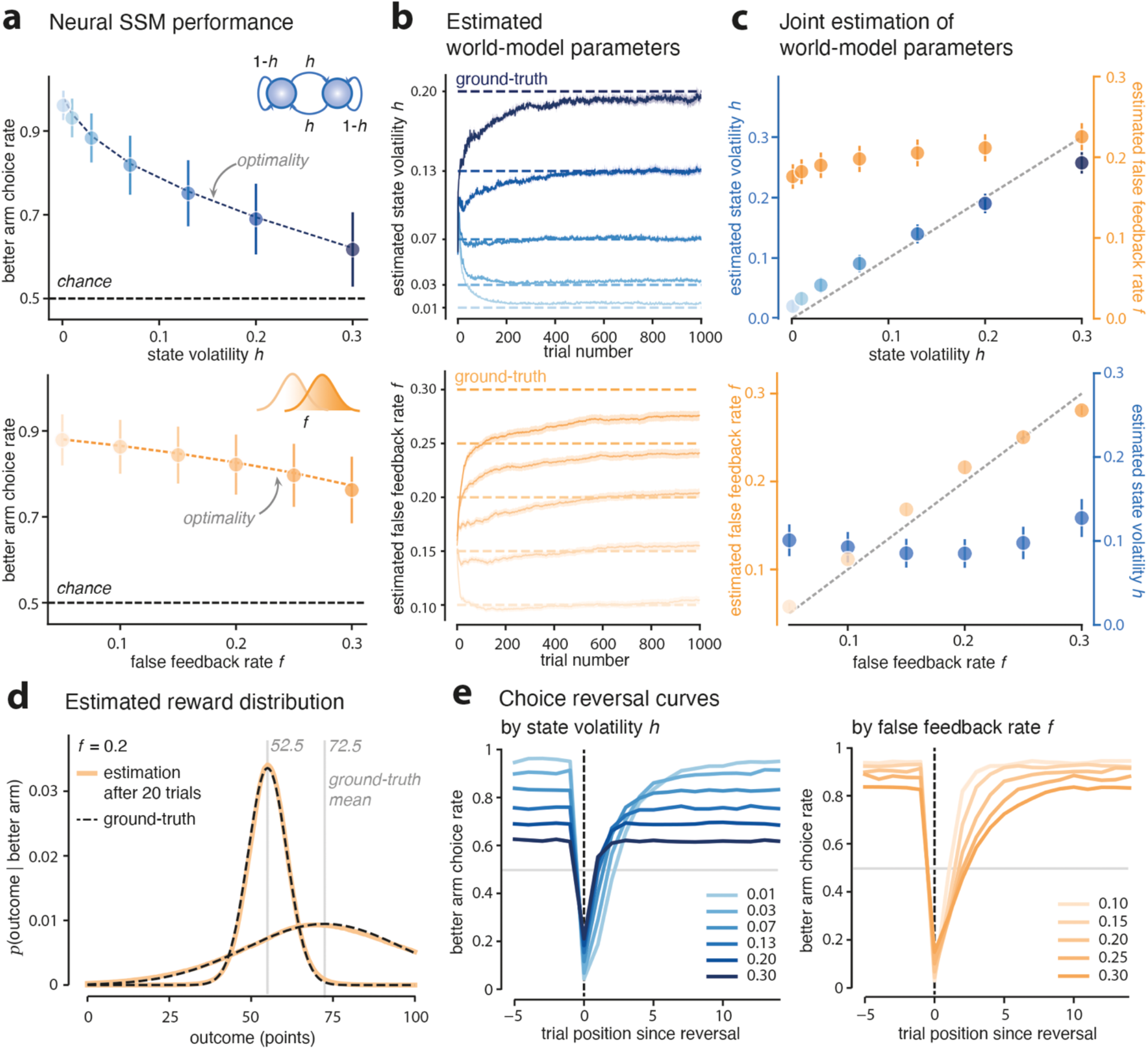
Neural state-space models learn generative world models in reversal learning. (**a**) Neural SSM performance. Top: proportion of correct choices (better arm choice rate) as a function of true state volatility *h*. Performance decreases as volatility increases, reflecting increased difficulty in tracking rapidly changing contingencies. Bottom: proportion of correct choices as a function of true false-feedback rate *f*. Larger *f* values increase the overlap between outcome distributions for the better and worse arms. Performance decreases as emission uncertainty increases. Solid lines indicate model performance; dashed grey lines indicate the performances of the optimal Bayesian model. (**b**) Estimated world-model parameters. Top: trial-by-trial estimates of state volatility for different true volatility levels (dashed lines). Bottom: trial-by-trial estimates of false-feedback rate for different true false-feedback levels. Estimates converge toward their respective groundtruth values over trials, demonstrating accurate recovery of both transition and false-feedback parameters. (**c**) Joint estimation of world-model parameters. Top: estimated state volatility and false-feedback as a function of true volatility. Bottom: estimated false-feedback rate and state volatility as a function of true false-feedback rate. (**d**) Estimated reward distribution. Example estimated emission distribution *p*(*y*_*t*_| *better arm*) after 20 trials (solid line), compared to the groundtruth distribution (dashed line). Neural SSMs recover the shape of outcome distributions, demonstrating flexible learning of non-parametric emission densities. (**e**) Reversal-aligned behavior. Left: choice accuracy aligned to reversal events for different volatility levels. Higher volatility produces faster post-reversal adaptation, consistent with larger inferred volatility states. Right: reversal-aligned performance for different false-feedback levels, showing that emission uncertainty affects asymptotic performance but does not induce the same change in learning speed as volatility.

At the level of latent parameter inference, neural SSMs accurately recover both sources of uncertainty (**Fig. 2b**). Trial-by-trial estimates of state volatility rapidly converge to values that closely match the true environmental volatility across conditions (**Fig. 2b**, top). Conversely, trial-by-trial estimates of the false-feedback rate converge toward true false-feedback levels when outcome reliability is manipulated (**Fig. 2b**, bottom). Critically, neural SSMs dissociate these two sources of uncertainty in their estimates (**Fig. 2c**). The estimated state volatility increases monotonically with the true environmental volatility while remaining largely insensitive to changes in outcome reliability (**Fig. 2c**, top), whereas the estimated false-feedback rate grows with the true false-feedback rate while remaining largely independent of environmental volatility (**Fig. 2c**, bottom). The emission network of neural SSMs accurately estimates not only the false-feedback rate, but also the shape of sampling distributions of rewards (**Fig. 2d**). The same trial-by-trial dynamics and parameter relationships are obtained with the optimal Bayesian model (**Supplementary Fig. 1**), showing that neural SSMs closely reproduce Bayes-optimal inference. Together, these results show that neural SSMs jointly recover world-model parameters, dissociate transition and emission uncertainty, and adjust learning rates appropriately after reversals (**Fig. 2e**), supporting amortized world-model learning.

We next examined **associative learning** using the **stimulus prediction task** – also referred to as the ‘weather prediction’ task^24–27^ – in which uncertainty arises from unknown and evolving associations between cues and stimuli rather than from changes in action-outcome contingencies (**Fig. 3a**). On each trial, a sequence of symbols *x* (cues) is sampled probabilistically conditioned on a stimulus *y* (red or green), and the agent must predict the stimulus identity from sampled symbols before observing the stimulus. Because cue-stimulus associations (e.g., *p*(*x* = star|*y* = red) = 0.8) can change unpredictably across time, successful performance in this task requires sustained learning and flexible updating of associative structure (i.e., conditional probabilities between cues and stimuli).

**Figure 3.**
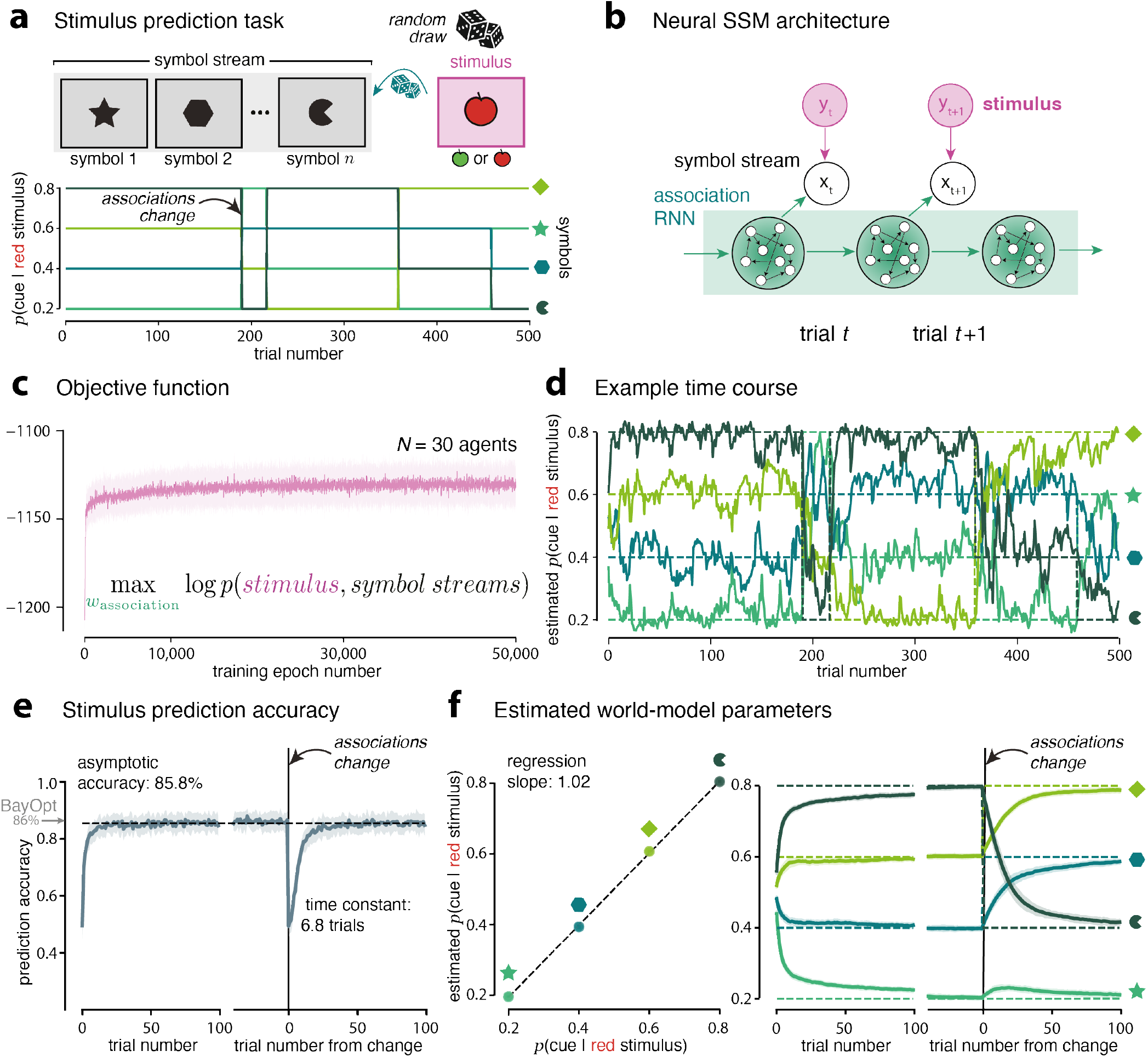
Neural state-space models learn generative world models in associative learning. (**a**) Stimulus prediction task. On each trial, a hidden stimulus *y* (red or green apple) generates a set of cues *x* (symbols) based on cue-stimulus associations. The agent must predict the stimulus from observed cues, before the true stimulus identity is revealed. Cue–stimulus associations are unknown and can change over time, requiring sustained learning of the associative structure. (**b**) Neural state-space model. A latent state encodes stimulus identity on each trial. A learned associative network outputs a four-dimensional vector representing stimulus-specific conditional probabilities *p*(*x*_*k*_| *y* = red). These probabilities are accumulated in log-odds space to compute the posterior belief over stimulus identity. (**c**) Training dynamics. Objective function (log-likelihood of true stimuli) as a function of training epochs, averaged across *N* = 30 independently trained neural SSMs, showing reliable convergence. (**d**) Trial-by-trial estimates of cue-stimulus associations. Example trajectories from a trained neural SSM showing sustained updating of associative structure as cue-stimulus associations change. (**e**) Stimulus prediction accuracy. Proportion of trials in which the hidden stimulus was correctly predicted, shown as a function of trial number. Performance is shown during initial learning and aligned to changes in cue–stimulus associations, showing adaptation following associative changes (BayOpt: optimal Bayesian model). (**f**) Estimated world-model parameters. Left: relationship between true cue-stimulus associations and the corresponding asymptotic estimates of neural SSMs. Right: time courses of learned cue-stimulus associations, demonstrating flexible updating of the estimated associative structure.

In this task, neural SSMs learned an **associative network** encoding cue-stimulus associations. The associative network estimated, for the stimulus *y* = red, the conditional probability that it indicated each cue *x*_*k*_, *p*(*x* = *x*_*k*_|*y* = red) (**Fig. 3b**). On each trial, these conditional probabilities were integrated across observed cues in log-odds space to compute the posterior belief over the stimulus identity, which constituted the prediction of the model. The associative network updated its estimates based on two inputs: the true stimulus identity observed at the end of the previous trial, and the counts of each cue in the previous trial, allowing it to track evolving cue-stimulus associations over time. Training neural SSMs by maximizing the likelihood of observed (true) stimuli led to reliable convergence across *N* = 30 neural SSMs trained independently (**Fig. 3c**).

An example time course of the estimated *p*(*x* = *x*_*k*_|*y* = red) for a trained neural SSM is shown in **Fig. 3d**. These trial-by-trial outputs of the associative network illustrate the amortized learning of cue-stimulus conditional probabilities over a single task realization. Prediction accuracy improved during initial learning and adapted rapidly following abrupt changes in cue-stimulus associations (**Fig. 3e**). Asymptotic performance reached 85.8% of accurate predictions, only 0.2% below the optimal Bayesian model. The associative network also accurately recovered the true associative structure. As shown in **Fig. 3f** (left), the relationship between true and estimated cue-stimulus associations had a slope of 1.02, indicating bias-free recovery of associative structure. A direct comparison of neural SSMs with the optimal Bayesian model shows similar prediction accuracy and parameter recovery, with only slightly faster adaptation to changes in the optimal Bayesian model (**Supplementary Fig. 2**). **Fig. 3f** (right) shows the time courses of inferred cue-stimulus associations, demonstrating flexible updating after changes. Together, these results show that neural SSMs accurately learn the associative structure of the task, and that the learnt parameters are sufficient to support near-optimal predictions by summing them across cues.

To further examine the generality of this framework, we trained neural SSMs to solve a more cognitively demanding task – a variant of the ‘mastermind’ task in which the agent must decipher a hidden combination from trial-to-trial feedback^10,28^. Again, trained neural SSMs achieved near-optimal choice accuracy and recovery of world-model parameters (**Supplementary Fig. 3**). These results indicate that the neural SSM framework extends beyond simple world models and remains effective in tasks involving a richer combinatorial structure. Taken together, our findings show that neural SSMs support amortized learning of structured world-model components in cognitive settings where classical Bayesian approaches require explicit model specification and computationally demanding inference schemes.

### Zero-shot generalization through re-use of learned world-model components

By design, neural SSMs introduce strong inductive biases for modular world-model structure, by allowing world-model components learned in one task to be reused and stacked in novel conditions that share properties with previously encountered ones. We therefore asked whether the learned transition, emission, and associative networks could be reused to solve tasks beyond the ones on which they were trained. Specifically, we examined whether previously learned world-model components could be recombined to solve a new task without any additional re-training.

To test this, we considered a category prediction task (**Fig. 4**) that combines **associative uncertainty** as in the stimulus prediction task (**Fig. 3**) with **emission uncertainty** as in the two-armed bandit task (**Fig. 1**). On each trial, a latent category (red or green) generates a sequence of cues (symbols) according to unknown cue-category associations (**Fig. 4a**). After observing the sequence of cues, the agent must predict the category, and subsequently observes a stimulus (colored apple) providing ambiguous feedback about the true category – by sampling colored stimuli from overlapping category-specific distributions. These overlapping stimulus distributions mirror the overlapping reward distributions associated with the better and worse arms of the two-armed bandit task (**Fig. 1**). Importantly, the true category is never revealed directly, requiring the agent to integrate the ambiguous evidence provided by cues with the ambiguous evidence provided by the observed stimulus to jointly learn the associative structure and the emission structure. We constructed neural SSMs by reusing previously learned modules rather than training neural SSMs from scratch. The **associative network** learned in the **stimulus prediction task** (**Fig. 3**) provides estimates of cue-category relationships, while the **emission network** learned in the **two-armed bandit task** (**Fig. 2**) learns the overlapping category-specific distributions of stimuli (**Fig. 4b**; see **Methods**).

**Figure 4.**
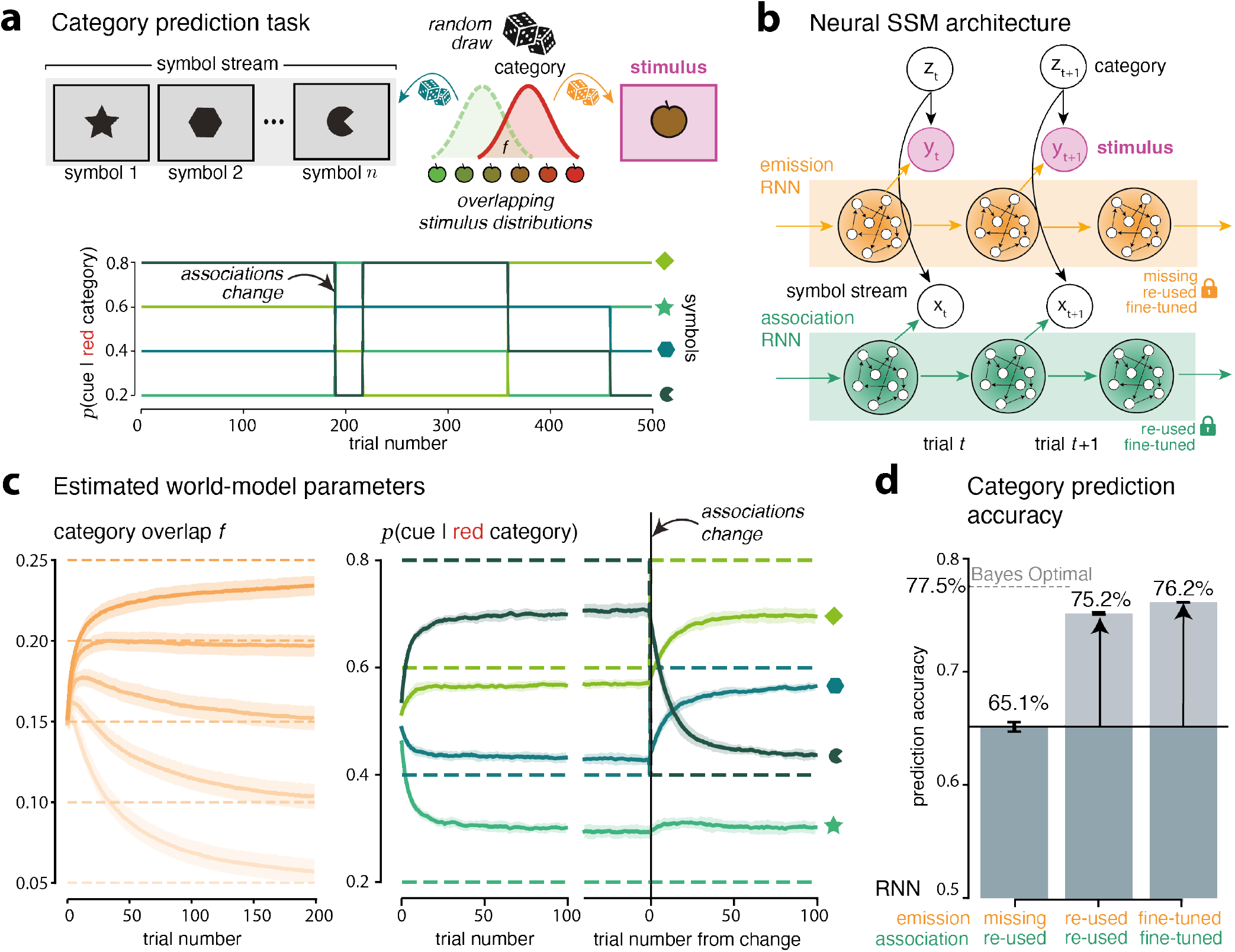
Zero-shot generalization through component re-use in category prediction. (**a**) Category prediction task. Top: on each trial, a latent category (red or green) generates a sequence of symbols (cues) according to cue-category associations that are unknown and change over time. The agent must predict the latent category from observed cues. After each choice, a continuous stimulus (colored apple) is drawn from the associated category-specific distribution, introducing emission uncertainty controlled by the category-overlap parameter *f*. Because the latent category is never revealed, learning must rely on ambiguous evidence provided by stimuli. Bottom: example trajectories illustrating trial-by-trial cue-category associations and un-signaled changes. (**b**) Neural SSM architecture with component re-use. A latent state *z* encodes category identity. As in the stimulus prediction task, the associative network outputs a four-dimensional vector representing stimulus-specific conditional probabilities *p*(*x*_*k*_| *y* = red). These values are accumulated across observed cues in log-odds space to compute the posterior belief over category identity. The emission network models category-specific probability distributions of stimuli. (**c**) Recovery of category overlap and cue-category associations. Estimated world-model parameters for neural SSMs in which the association and emission modules were re-used. Left: trial-by-trial estimates of the category-overlap parameter *f*, reflecting emission uncertainty. Right: time courses of estimated cue-category associations, demonstrating flexible updating of estimated associative structure. (**d**) Category prediction accuracy across neural SSMs with different configurations. Left: only the associative module is re-used. Ambiguous stimuli are therefore considered as certain evidence and provided directly to the associative network, as in the stimulus prediction task. Center: both the associative and emission networks are re-used without fine-tuning (as in **c**). Right: networks are additionally fine-tuned on the category prediction task. Re-used networks already yield prediction accuracy levels close to optimal Bayesian inference (within 2.3_*k*_), which further improve with additional fine-tuning (within 1.3_*k*_).

Although the emission and association networks were trained independently on different tasks, stacking them in the same neural SSM enabled accurate inference over both cue-category associations and emission uncertainty (**Fig. 4c**), achieving performance within 2.3% of optimal Bayesian inference (**Fig. 4d**). Trial-by-trial estimates showed accurate recovery of both emission uncertainty, quantified by the category-overlap parameter *f*, and cue-category associations as they evolved over trials (**Fig. 4c**; see **Supplementary Fig. 4** for the optimal Bayesian model). Category prediction accuracy depended strongly on the configuration of the neural SSM (**Fig. 4d**). Re-using only the associative network and treating the ambiguous stimulus as certain feedback (i.e., *f* = 0) led to substantially reduced performance (**Fig. 4d** and **Supplementary Fig. 4**). Fine-tuning the re-used networks to the category prediction task produced additional – but modest – improvements (reaching 1.3% of optimal Bayesian inference). This result indicates that the re-used associative and emission networks can already learn most of the world-model structure required for solving the category prediction, task while allowing for further task-specific refinements.

For comparison, we also trained neural SSMs from scratch on the category prediction task (**Supplementary Fig. 5**). Entropy regularization was required during training for neural SSMs to achieve prediction accuracy levels comparable to that of neural SSMs with re-used components (**Supplementary Fig. 5**; see **Methods**). However, even with entropy-regularized training, learning from scratch in the category prediction task proceeded substantially slower than training the emission network in the two-armed bandit task and the associative network in the stimulus prediction task, and re-using them to solve the category prediction task without any additional training (**Supplementary Fig. 5**). Moreover, although neural SSMs trained from scratch eventually reached near-optimal prediction accuracy, it recovered the underlying cue-category associations substantially less accurately (**Supplementary Fig. 5**). Together, these results show that the re-use of world-model components in neural SSMs not only accelerates learning but also promotes representations that more faithfully capture the world-model structure of the novel task on which they are tested.

To further test the generality of component re-use in neural SSMs, we considered a history-dependent variant of the category prediction task in which the latent category evolves over time – with a transition structure mirroring changes in the better and worse arms of the two-armed bandit task (**Fig. 5a**). In addition to learning cue-category associations and interpreting ambiguous stimuli, agents must now exploit temporal dependencies in the category identity across trials – governed by a state volatility parameter *h*. This task therefore combines the three sources of uncertainty encountered in previous tasks: **associative uncertainty, emission uncertainty**, and **transition uncertainty**. We constructed neural SSMs by re-using the three networks trained in previous tasks without any additional re-training: the transition and the emission networks from the two-armed bandit task, and the associative network from the stimulus prediction task (**Fig. 5b**). These components were combined to perform inference over the latent category while integrating information from presented cues and ambiguous stimuli sampled from the latent category.

**Figure 5.**
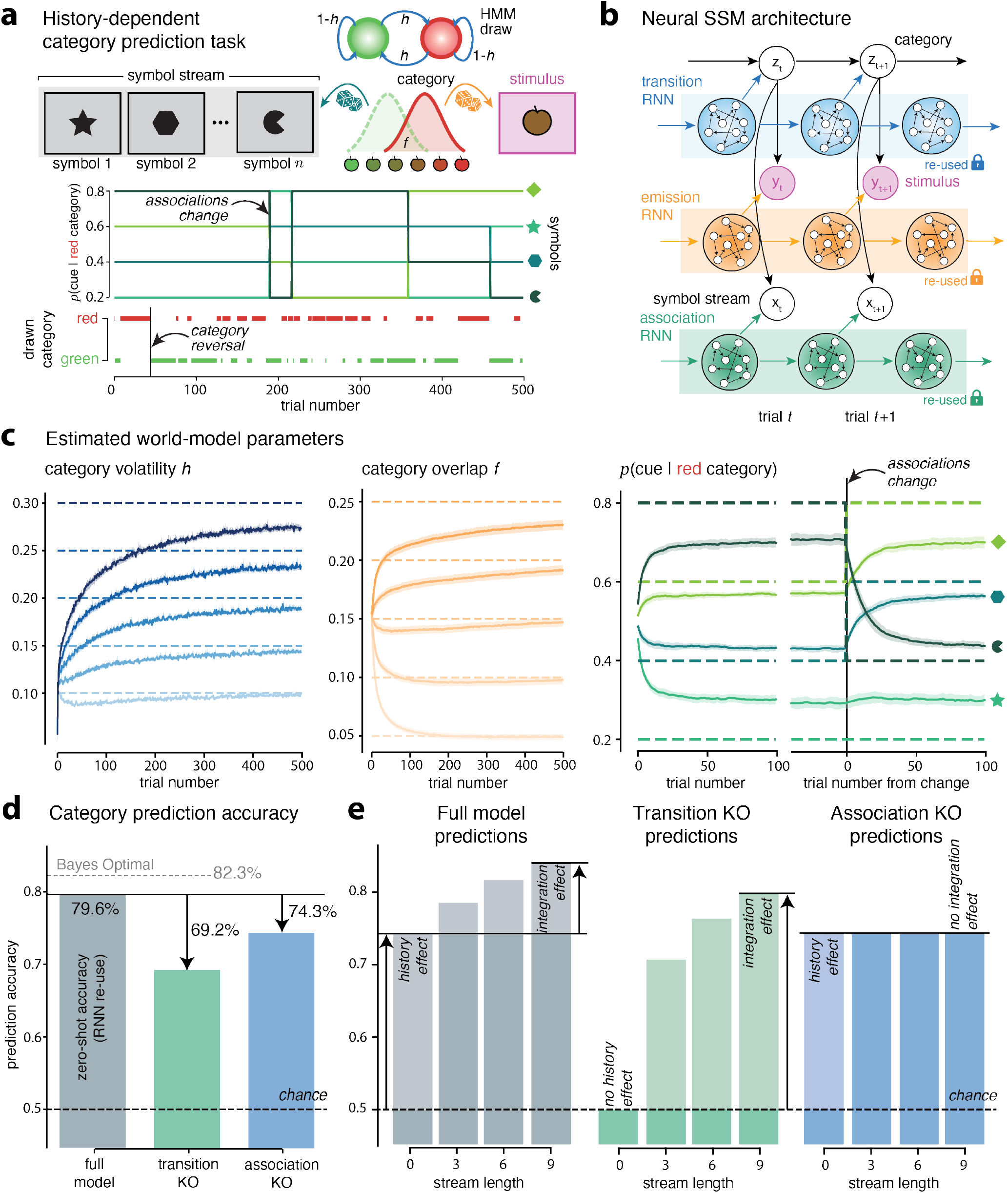
Zero-shot generalization through component re-use in history-dependent category prediction. (**a**) History-dependent category prediction task. Top: on each trial, a latent category (red or green) evolves according to a two-state Markov process with volatility *h*, introducing transition uncertainty. All other aspects of the task are identical to the category prediction task. Bottom: example trajectories illustrating trial-by-trial cue-category associations and un-signaled changes (upper panel), and the Markovian evolution of the latent category across trials (lower panel). (**b**) Neural SSM architecture with component re-use. Neural SSMs combine the three components learned in previous tasks without any additional re-training: a transition network estimating category volatility, an emission network modeling category-specific stimulus distributions, and an associative network encoding cue-category associations. A latent state z represents category identity on each trial. The associative and emission networks are re-used as in the category prediction task. The transition module captures temporal correlations in category identity across trials. (**c**) Estimated world-model parameters. Left: trialby-trial estimates of the category volatility parameter *h*. Middle: trial-by-trial estimates of the category-overlap parameter *f*. Right: time courses of estimated cue-category associations. (**d**) Category prediction accuracy across neural SSMs with different configurations. The full neural SSM achieves a performance of 79.6%, close to optimal Bayesian inference (82.3%). Knocking out either the transition network or the associative network significantly reduces accuracy (69.2% and 74.3%, respectively). (**e**) Selective impact of component knock-out on history and integration effects. Prediction accuracy is shown as a function of symbol-stream length for full neural SSMs and knocked-out variants. Full neural SSMs exhibit history and integration effects – benefitting from temporal correlations across trials and from additional cues within each trial. Neural SSMs lacking a transition network show zero history effect, whereas neural SSMs lacking an associative network show zero integration effect (no increase in prediction accuracy with more cues).

Despite never being trained on this complex task, recombined neural SSMs successfully recovered all components of the world-model structure. Trial-by-trial estimates tracked the state volatility parameter governing category transitions, the category-overlap parameter controlling category-specific stimulus distributions, and the evolving cue-category associations (**Fig. 5c**; see **Supplementary Fig. 6** for the optimal Bayesian model). These estimates demonstrate that the re-used components collectively form a coherent inference system capable of simultaneously handling multiple sources of uncertainty in the absence of any task-specific re-training. Component re-use supported zero-shot behavioral performance, achieving 79.6% prediction accuracy (**Fig. 5d**), within 2.7% from optimal Bayesian inference. To assess the contribution of each component, we selectively knocked-out (eliminated) individual components from recombined neural SSMs. Eliminating either the **transition network** or the **associative network** produced substantial drops in prediction accuracy, indicating that accurate predictions rely both on long-term temporal correlations of the latent category across trials and on short-term predictive information provided by the sequence of cues within each trial. Further analyses revealed distinct functional roles for the reused modules (**Fig. 5e**). Full neural SSMs exhibited both a **history eJect** – benefitting from temporal correlations across trials – and an **integration eJect** – benefitting from more cues within each trial. Knocking out the transition network eliminated the history effect, whereas knocking out the associative network abolished the integration effect. These comparisons show that each re-used component contributes complementary computations which, together, support accurate inference of the latent category in the more complex history-dependent environment.

Taken together, this pattern of results demonstrates that learned world-model components can be recombined in neural SSMs to support near-optimal inference in environments that differ from those encountered during training. Rather than jointly re-learning all components from scratch for each new task, neural SSMs can leverage structure acquired separately in previous tasks to solve new tasks involving multiple sources of uncertainty. This framework provides a modular implementation of the Bayesian Brain capable of flexible component re-use and curriculum-like learning across tasks of increasing complexity.

### Behavior signatures of component re-use in human reversal learning

The results obtained so far show that neural SSMs can recombine components of a generative world model learnt across previous tasks to solve a new task that requires (some of) the same components. If similar principles govern human learning, then individuals should also re-use the same components to solve different tasks that share the same generative structure, even when the tasks appear superficially different. To test this prediction, we designed two reversal learning tasks with superficially dissimilar characteristics but identical latent transition structure (**Fig. 6a**). In the **bandit task**, participants chose between two symbols (arms) and received a continuously-valued reward (outcome) sampled from the probability distribution associated with the chosen symbol – an instrumental learning paradigm. In the **apples task**, the same participants observed streams of colored apples sampled from one of two possible sources (each associated with a probability distribution), and reported after each apple the currently sampled source – a visual categorization paradigm. Despite the many superficial differences between the tasks, both were governed by the same transition structure: a latent binary state determining the better arm (in the bandit task) or the currently sampled source of apples (in the apples task). This latent state evolved according to a two-state Markov process with state volatility *h*, introducing the same transition uncertainty in both tasks (**Fig. 6b**; see **Methods**).

**Figure 6.**
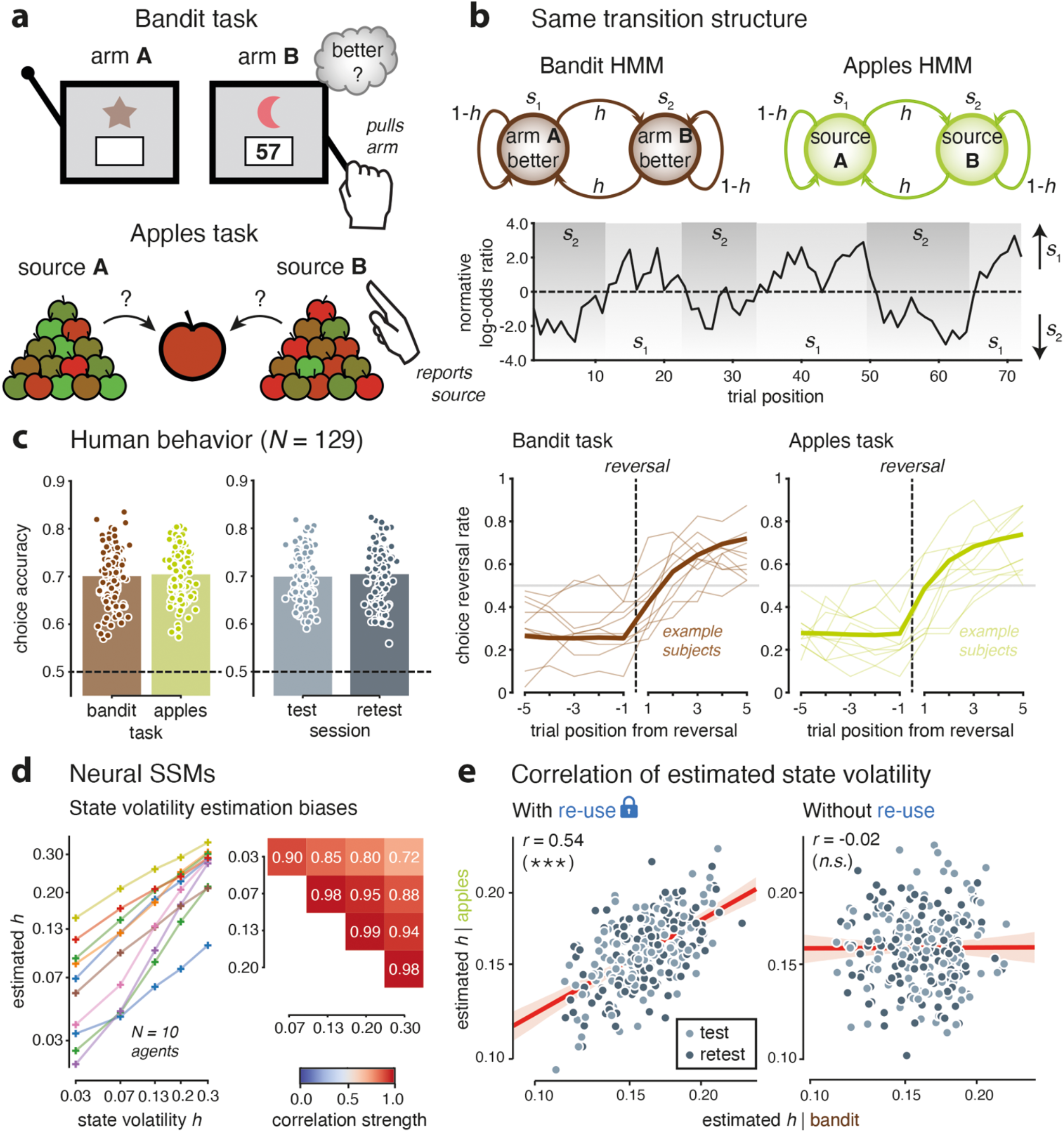
Signatures of component re-use across superficially dissimilar tasks (**a**) Task description. Top: bandit task. On each trial, the agent selects between arms (A or B) and receives a continuous-valued reward. Bottom: apples task. On each trial, the observer reports the source (A or B) from which apples are currently being sampled. (**b**) Same transition structure between tasks. Left: in the bandit task, a latent binary state determines which arm is currently better. This latent state evolves according to a Markov process with state volatility *h*. Right: in the apples task, a latent binary state determines which source of apples is currently sampled. This latent state evolves according to the same Markov process as in the bandit task, with the same state volatility *h*. Bottom: example transition instance, showing transitions between states *s*^1^ and *s*^2^, and the normative log-odds ratio associated with rewards (bandit task) or apples (apples task). (**c**) Human behavior (*N* = 129 participants). Left: same choice accuracy across tasks (left), and across sessions (right). Right: similar choice reversal curves in the bandit (left) and apples (right) tasks. (**d**) State volatility estimation biases in neural SSMs. Left: relation between the true state volatility *h* (x-axis) and the estimated state volatility *h* (y-axis) for *N* = 10 neural SSMs trained independently in the same conditions. All correlation strengths are above 0.7. Right: correlation matrix of estimated state volatilities across levels of true state volatility. (**e**) Between-task correlation of estimated volatility in neural SSMs (x-axis: in bandit task; y-axis: in apples task). Left: significant between-task correlation with re-use of the same transition network in the two tasks. Right: zero between-task correlation without component re-use – i.e., different transition networks in the two tasks. Three stars indicate a significant effect at *p* < 0.001, n.s. a non-significant effect.

Each participant (*N* = 129 after attrition; see **Methods**) performed the bandit and apples tasks in separate runs, and completed the same experiment again two weeks later (T_1_: test session, T_2_: retest session). Participants performed both tasks well above chance level, and achieved comparable levels of choice accuracy across tasks and across sessions (**Fig. 6c**; paired *t*-test, between tasks: *p* > 0.2, between sessions: *p* > 0.2). Examining behavioral dynamics revealed clear adaptation to changes in the latent state: following reversals, participants gradually adjusted their choices, producing similar reversal-aligned learning curves in both tasks (**Fig. 6c**). These results show that participants successfully tracked changes in the latent state from ambiguous action outcomes (rewards in the bandit task) and from ambiguous observations (colored apples in the apples task). The behavioral data also showed large individual differences between participants (**Fig. 6c**), which we leveraged to identify signatures of component re-use across the two tasks.

First, we turned to neural SSMs trained in the **bandit task** to assess their differences after training on different instances of the same task (**Fig. 6d**). Across *N* = 10 trained neural SSMs – selected to span the full range of estimated state volatility (based on average estimates) – we found that neural SSMs differ reliably in their estimation of state volatility (measured at the end of each bandit task instance) when tested across different levels of state volatility. Some neural SSMs overestimate state volatility, whereas others underestimate the same world-model parameter (**Fig. 6d**, left). The reliability of these individual differences in trained neural SSMs was evident when plotting the correlation of state volatility estimates across different levels of true state volatility (**Fig. 6d**, right). These results indicate that the transition components differ reliably across neural SSMs trained on different instances of the same task.

We then constructed neural SSMs for the **apples task** by re-using the transition and emission networks learnt in the bandit task (**Fig. 1b**) – by modifying the inference according to the characteristics of the apples task without any re-training of the world-model components. We compared neural SSMs which re-used the same transition component across the two tasks – by using the transition network of the same neural SSM in the two tasks – with neural SSMs which used different transition components across the two tasks – by using the transition networks of different neural SSMs in the two tasks. We found that the average estimated state volatility correlates strongly across tasks (*r* = 0.54, *p* < 0.001) for neural SSMs with component re-use (i.e., the same transition network across tasks), but does not correlate across tasks (*r* = -0.02, *p* > 0.5) for neural SSMs without component reuse (i.e., different transition networks across tasks; **Fig. 6e**). This result shows that the correlation between estimated state volatility across the bandit and apples tasks is diagnostic of component reuse across the two tasks.

To look for the same signature in human data, we fitted each participant’s behavior in each task (bandit, apples) and each session (test, retest) with a previously validated **hidden-state inference model**^29–32^ with two free parameters: 1. the **subjective volatility** *h* of the latent state (the better arm in the bandit task, the sampled source in the apples task), and 2. the **temperature** *τ* of the softmax policy used to choose – between arms in the bandit task, between sources in the apples task (see **Methods**). To account for the superficial differences between the two tasks, we additionally modeled task-specific biases that have been described in similar conditions^33–36^. Specifically, we modeled a **subjective utility function** accounting for loss aversion when rewards fall below a reference point in the bandit task, and a **category evidence function** accounting for category bias and attraction effects in the apples task. (**Supplementary Fig. 7**; see **Methods**). We first verified that individual differences in state volatility *h* and policy temperature *τ* measured in each task correlated across sessions (**Fig. 7a**; state volatility: *r* = 0.76, *p* < 0.001; policy temperature: *r* = 0.55, *p* < 0.001). This means that both parameters reflect trait-like strategies in participants which remain stable across weeks.

**Figure 7.**
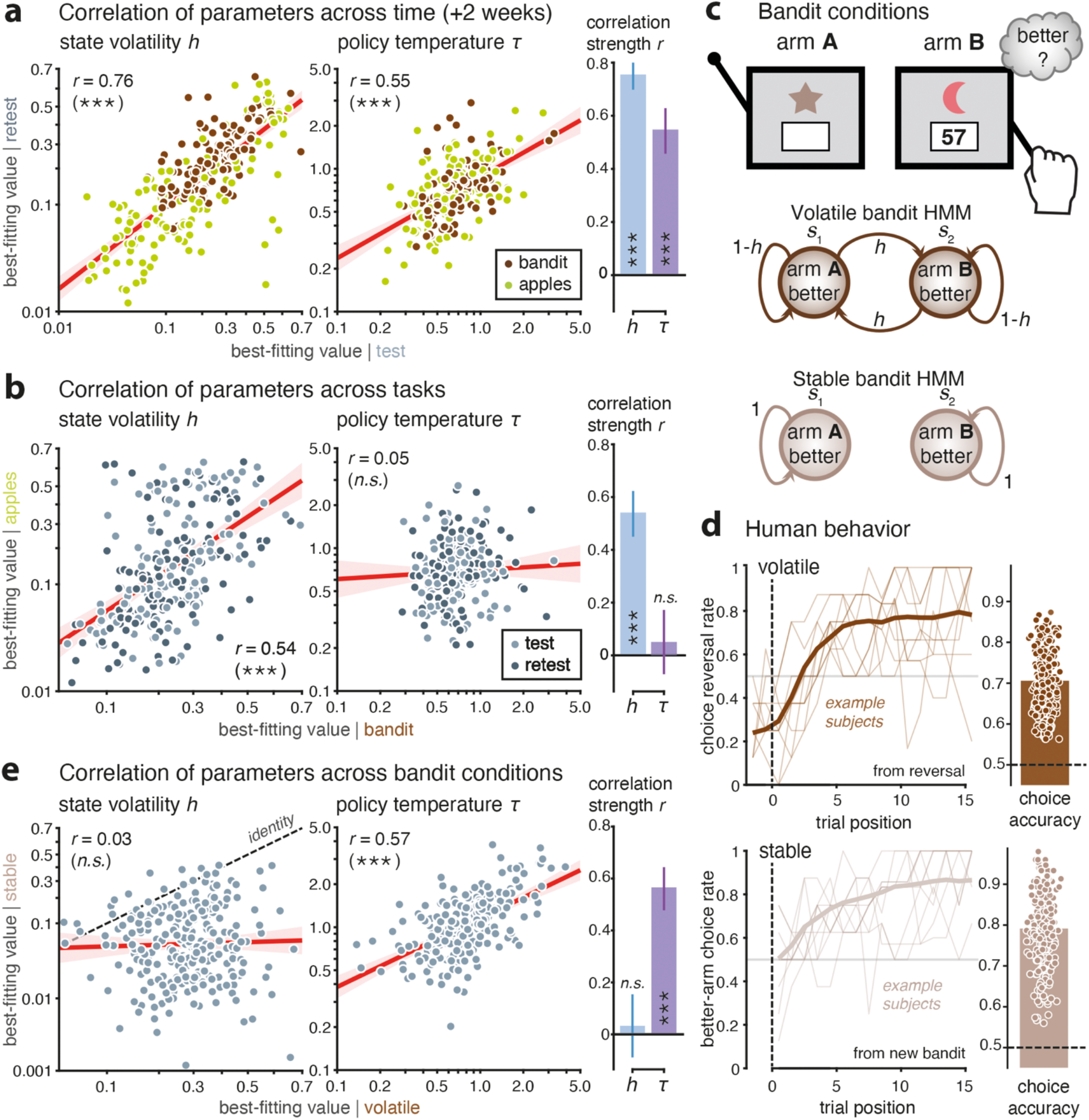
Evidence of component re-use in humans. (**a**) Correlation of parameters across time (x-axis: test session; y-axis: retest session). Left: strong correlation of state volatility *h* (left) and of policy temperature τ (right) across time in both tasks (brown dots: bandit task; green dots: apples task). Right: correlation strength across time for *h* and τ. Error bars indicate 95% CI. (**b**) Parameter correlation between tasks (x-axis: bandit; y-axis: apples). Left: strong correlation of state volatility *h* (left) but zero correlation of policy temperature τ (right) between tasks (lighter dots: test; darker dots: retest). Right: correlation strength between tasks for *h* and τ. (**c**) Description of bandit task conditions. Top: on each trial, the agent selects between arms and receives a continuous-valued reward. Bottom: transition structures in the volatile (top) and stable (bottom) bandit conditions. A latent state determines the better arm. In the volatile condition, the latent state evolves according to a Markov process with state volatility *h*. In the stable condition, the latent state does not change over the course of a bandit game. (**d**) Human behavior (N = 260 participants). Top: choice reversal curve in the volatile condition (left) and associated choice accuracy (right). Bottom: better-arm choice rate in the stable condition (left) and associated choice accuracy (right). (**e**) Parameter correlation between bandit conditions (x-axis: volatile; y-axis: stable). Left: zero correlation of state volatility *h* (left) but strong correlation of policy temperature τ (right) between conditions. Right: correlation strength between conditions for *h* and τ. Three stars indicate a significant effect at p < 0.001, n.s. a non-significant effect.

We then plotted each model parameter in the apples task against the same parameter in the bandit task (**Fig. 7b**) – our diagnostic signature of component re-use (**Fig. 6e**). We found that individual differences in subjective state volatility *h* correlate reliably across tasks (*r* = 0.54, *p* < 0.001) – as observed in neural SSMs with re-use of the same transition component across tasks. By contrast, individual differences in policy temperature did not correlate across tasks (*r* = 0.05, *p* > 0.2). This absence of correlation is consistent with the notion that, unlike the transition structure which is task-general, the decision policy is task-specific. In the bandit task, it corresponds to an instrumental ‘sampling’ policy governed by a trade-off between exploration and reward maximization^37–40^. By contrast, the decision policy used in the apples tasks reflects a non-instrumental ‘reporting’ policy which maps inferred beliefs to response categories. Parameter recovery analyses confirmed that these estimates are reliable and interpretable at the individual level (**Supplementary Fig. 7**). We further validated that the correlation of state volatility *h* across tasks does not arise from the testing of participants on identical transition instances between the two tasks (**Supplementary Fig. 8**; identical transition instances: *r* = 0.48, *p* < 0.001; different transition instances: *r* = 0.50, *p* < 0.001), and that individual differences in state volatility *h* are associated with diagnostic changes in reversal-aligned learning curves (**Supplementary Fig. 8**). We also confirmed that individual differences in state volatility *h* do not reflect differences in repetition bias (**Supplementary Fig. 9-10**), by showing that participants’ tendency to repeat their previous choice depends on the match between their previous choice and their prior belief (estimated through particle filtering; see **Methods**) – and more strongly so for participants with low subjective state volatility *h*.

The strong correlation in state volatility between the bandit and apples tasks suggests that humans re-use the same transition component to track the latent state in the two tasks – the better arm in the bandit task, the sampled source in the apples task – despite their glaring superficial differences. But does component re-use in human learning depend on context? The framework predicts that humans should not re-use the same transition component across tasks that do not share the same transition structure, even if they are superficially similar. To test this hypothesis, we examined component reuse in a published human behavior dataset^41^ where tested participants (*N* = 260 after attrition; see **Methods**) played, across different runs, two conditions of a bandit task: a **volatile condition** where arm values changed across time (as in the bandit task used in **Fig. 6**), and a **stable condition** where arm values had constant means across time (**Fig. 7c**). Unlike the previous paradigm where bandit and apples tasks were superficially dissimilar but shared the same transition structure, the two bandit conditions are superficially similar – both were described to participants as two-armed slot machine games – but they do not share the same transition structure – the better arm remained the same throughout each game in the stable condition. Human behavior showed characteristic learning curves in the two conditions (**Fig. 7d**), again with large individual differences in learning across tested participants.

Because the volatile and stable conditions do not share the same transition structure, we predicted that subjective state volatility *h* estimated from participants’ behavior should not correlate across bandit conditions. And indeed, this diagnostic signature of component re-use was not present in this dataset when comparing stable and volatile bandit conditions (**Fig. 7e**; *r* = 0.03, *p* > 0.5). By contrast, the best-fitting policy temperature *τ* was found to correlate very strongly between the two bandit conditions (*r* = 0.57, *p* < 0.001). This strong correlation is consistent with the notion that participants can re-use the same ‘explore-exploit’ choice policy across the stable and volatile conditions of the bandit task. Together, these findings provide further support for component re-use in human learning: humans appear to re-use the same transition component across superficially dissimilar tasks sharing the same latent transition structure, but not across conditions of the same task that rely on different transition structures.

## Discussion

Existing implementations of the Bayesian Brain hypothesis pre-specify both the form of the generative model and the inference algorithm^1,10–12^. Neither is learned, leaving little room for meta-learning, structural discovery, or flexible adaptation across tasks. We have shown that neural SSMs overcome these longstanding limitations by jointly learning both the world model and the inference dynamics. Unlike rigid state-space models, neural SSMs can discover functionally meaningful latent parameters that support prediction and adaptation^42,43^. In other words, we did not enforce that the transition network should learn to estimate volatility throughout training. Instead, neural SSMs are trained to best predict observed outcomes (e.g., rewards in the bandit task, colored stimuli in the stimulus prediction task). This Bayesian objective creates an **inductive bias** on the neural SSM architecture, which leads to a world-model component specialized in estimating a volatility-like parameter of the world model.

Inference over world-model parameters in neural SSMs is not carried out by a separate algorithm, but is implicitly implemented by the learned dynamics of the model, an instance of **amortized inference**^16,17^. In the four tasks considered here, performing Bayesian inference in the optimal generative model requires iterative, task-specific procedures. Rather than relying on such iterative variational or sampling-based procedures for parameter inference, neural SSMs compile inference into neural dynamics learned through experience. This results in orders-of-magnitude reductions in computation time (**Supplementary Table 1**). Amortized inference provides a computationally efficient and biologically plausible alternative to heavier inference schemes. Rather than asking how neural circuits implement specific inference algorithms such as sampling or particle filtering^44–47^, the key question becomes how neural circuits acquire dynamics that support accurate prediction and belief updating through learning.

The **fast-slow separation** in neural SSMs provides a concrete computational account of how humans might combine rapid inference with slower learning of re-usable components. At fast time scales, neural SSMs perform trial-by-trial belief updating through amortized neural dynamics, allowing them to continuously adapt to new observations signaling changes in world-model parameters. At slow time scales, the estimation of world-model parameters is itself learned across experience, promoting the acquisition of modular components of a world model that can be re-used across tasks that share the same (or similar) world models. This slow process can be described as a form of **metalearning**, through which neural SSMs acquire structural components of an internal model of the world that can be used to learn in different tasks (including the four tested in this study). Unlike end-to-end recurrent neural networks (RNNs) – where representations are entangled and task-specific^48,49^ – the explicit modularity of neural SSMs enables the separation and recombination of modular components, supporting generalization across tasks with superficial differences but identical latent structure. This computational re-use is consistent with the idea that the brain re-deploys shared computational motifs across cognitive tasks with compositional structure^19,20^.

Here, we implemented neural SSMs using RNNs, which provide a natural architecture for sequential data and offer a biologically plausible mechanism for maintaining and updating internal states over time. This makes RNNs well suited to implement amortized inference through dynamical state updates. However, the framework itself is not tied to a specific neural architecture. In principle, any **function approximator** capable of capturing temporal dependencies could be used to parameterize the different world-model components. For instance, transformer-based architectures^50^ – with their flexible attention mechanisms and capacity to capture long-range dependencies – could provide an alternative implementation of neural SSMs, with potentially richer representations in more complex real-life settings. Thus, the key contribution of the framework lies not in the choice of RNNs per se, but in the combination of learned generative structure and amortized inference – which can be instantiated across a broad class of neural architectures.

We have identified human signatures of computational re-use across superficially dissimilar tasks usually theorized in terms of vastly different **algorithmic models**: reinforcement learning (RL) models in bandit tasks^37,38,51,52^, drift-diffusion models in perceptual decision-making tasks^53–55^. Here, we have designed the bandit and apples tasks such that they are perfectly matched in terms of their transition structure. This perfect match allowed for the same hidden-state inference model^29–32^ to be applied to track the latent state of the two tasks – the better arm in the bandit task, the sampled source in the apples task. Equipped with these transition-matched tasks, we could then examine how the same individuals play both of them, fit the hidden-state inference model to each individual’s behavior in each task, and test whether some aspects of their strategies are shared between the two tasks despite their glaring superficial differences.

To identify computational re-use, we leveraged **individual differences** in behavior in the two tasks across hundreds of tested participants. Based on recent findings^56,57^, we reasoned that individual differences in each task are not random, but reflect genuine differences in the cognitive strategies deployed to solve both tasks. As predicted, individual differences in subjective volatility and in policy temperature showed strong correlations across time – two weeks in our study. These temporal correlations indicate that the slow learning of world-model components occurs across time scales much slower than weeks. Our framework assumes that transition, emission and associative components are present before starting the task, and acquired gradually by experience throughout development – or even innate in the wiring of neural circuits^58^.

Computational re-use in neural SSMs predicted selective correlations in volatility estimates across tasks using the same transition structure – something which had not been tested in humans to our knowledge. We observed this between-task correlation in human behavior, which suggests that individuals re-use some world-model components and construct a **mental graph** to solve a cognitive task. Indeed, individuals appeared to re-use the same transition component across different yet transition-matched tasks in the first dataset, and re-use the same decision policy across different bandit conditions in the second dataset. An important open question for future research is how humans build these mental graphs and converge on the re-use of specific world-model components in a context-dependent fashion^9,59^.

Our framework also generates a set of concrete and testable **neural predictions**. First, the brain should compute **state prediction errors** defined with respect to internal world models, rather than only reward prediction errors (RPEs). State prediction errors have already been theorized and reported by recent work in striatal and cortical circuits^60–63^. Second, different world-model components may be implemented by partially **dissociable neural substrates**. In other words, the brain may decompose inference into distinct processes – such as tracking changes in latent structure or learning associations – that are supported by separable but interacting neural circuits – like the transition and associative networks in neural SSMs – and can be flexibly recombined across tasks. For example, the estimation of state volatility and the learning of cue-stimulus associations may rely on partially distinct neural substrates, with the former involving medial frontal regions^12,52,64^ and the latter medial temporal regions^25,65,66^. Third, tasks sharing the same underlying generative structure should engage **overlapping belief update signals**, even when they differ in several other characteristics. In particular, the bandit and apples tasks should trigger similar belief update signals – in response to rewards in the bandit task, and to colored apples in the apples task – reflecting shared computations over latent structure across superficially dissimilar task environments.

This **augmented Bayesian Brain** framework provides a unified perspective on probabilistic reasoning, reward-guided learning, and compositional tasks as expressions of shared world-model components. We have provided behavioral evidence of world-model component sharing across tasks in humans. The framework makes testable neural predictions for future research in terms of between-task transfer of computations, including shared latent trajectories and overlapping dynamical motifs. Together, our findings suggest that flexible cognition emerges from learned, re-usable world-model components that can be combined across tasks and implemented through amortized neural dynamics.

## Methods

### Two-armed bandit task

On each trial *t*, the agent chose between two arms. A latent binary state *z*_*t*_ ∈ {0, 1} encoded which symbol was currently the better option (the higher-rewarding one). The latent state evolved over trials according to a Markov process with state volatility (reversal probability) *h*:

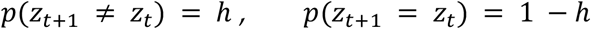

Thus, *h* governed the transition uncertainty, determining how frequently reversals occurred. *h* is sampled from an exponential distribution:

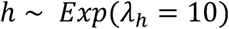

truncated to the interval [0, 0.4]. After a choice *a*_*t*_∈ {0, 1}, the agent received a continuous-valued outcome *y*_*t*_∈ [−1, 1]. Outcomes were drawn from emission distributions:

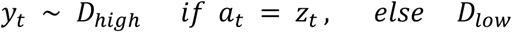

where *D*_*high*_ and *D*_*low*_ were overlapping reward distributions.

The high-reward emission distribution *D*_*high*_ was defined as a Gaussian:

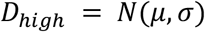

truncated to the interval [−1, 1]. The mean *µ* was sampled for each environment from an exponential distribution

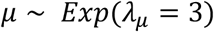

and truncated between [0, 1].

The standard deviation *σ* was determined by an emission-uncertainty parameter, referred to as the false-feedback rate *f*, sampled from:

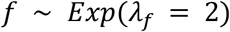

and truncated between [0, 0.4].

The false-feedback rate *f* was defined as the probability mass of the high-reward distribution below zero:

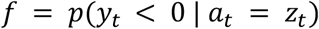

The low-reward distribution *D*_*low*_ was defined as the reflection of *D*_*high*_ around zero.

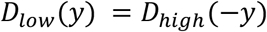

ensuring symmetric reward structure around zero. Larger values of *f* increased the overlap between reward distributions and reduced outcome reliability.

### Neural SSM architecture in the two-armed bandit task

The neural state-space model (neural SSM) is composed of a discrete latent Markov state encoding the currently better option and recurrent neural modules that learned to estimate transition and emission parameters from experience.

The latent state *z*_*t*_encoded which arm was currently better. The agent maintained a joint likelihood *p*(*z*_*t*_, *y*_1:*t*_ | *a*_1:*t*_) and a posterior belief *p*(*z*_*t*_| *y*_1:(*t*-1)_, *a*_1:(*t*-1)_) updated with likelihood recursion^23^; *a*_*t*_ and *y*_*t*_ are respectively the chosen action and the observed outcome at trial t.

At each trial, the agent selected the action corresponding to the arm with the highest posterior probability of being the better option:

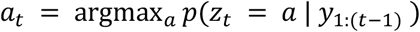

The emission distribution *p*(*y*_*t*_| *a*_*t*_= *z*_*t*_) was parameterized by a gated recurrent unit (GRU) network *p*_θ_ (32 units) that learned the full outcome density from trial history. To be noted that the emission distribution was produced by a recurrent neural network whose hidden state evolved across trials, such that it depended at each trial on the history of past observations through this internal state. At each trial, its hidden state was updated using two inputs, the current posterior belief over the selected state *p*(*z*_*t*_= *a*_*t*_| *y*_1:(*t*-1)_) and the observed outcome *y*_*t*_.

Rather than assuming a parametric form (e.g., Gaussian), the outcome space [−1, 1] was discretized into bins with resolution 0.01, yielding a grid of 201 bins. The emission network produced a probability mass function over this discretized grid at each trial. This flexible representation allowed the model to learn arbitrary outcome distributions without imposing assumptions on their shape. The GRU hidden state was mapped through a linear layer of dimension 201 to produce unnormalized values over the discretized outcome grid. These values were passed through a sigmoid nonlinearity and subsequently normalized to obtain a valid probability distribution over outcomes.

The emission distribution for the alternative hypothesis *p*(*y*_*t*_| *a*_*t*_≠ *z*_*t*_) was defined as the reflection of the learned distribution around zero: *p*(*y*_*t*_| *a*_*t*_≠ *z*_*t*_) = *p*_3_(−*y*_*t*_), ensuring a symmetric structure between the higher- and lower-rewarding arms.

The transition network was also implemented as a GRU network *p*_ϕ_ (32 units). At each trial t, the transition module received as input a scalar evidence signal derived from the current emission model. Specifically, after observing outcome *y*_*t*_ following action *a*_*t*_, we computed the log-likelihood ratio of the observed outcome between the hypothesis that the chosen arm matched the latent better state and the alternative hypothesis:

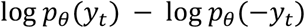

This input summarizes how strongly the observed outcome supports the belief that the chosen arm was indeed the better arm. The GRU network updated its hidden state based on this input, and the current reversal probability *h*_*t*_ was obtained by linearly projecting the hidden state to a scalar and applying a sigmoid nonlinearity.

### Neural SSM training in the two-armed bandit task

Model parameters, including emission parameters *θ* and transition parameters *ϕ*, were optimized by maximizing the log-likelihood of observed outcomes under the generative model:

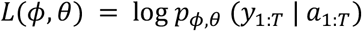

where *T* denotes the number of trials, *y*_1:*T*_ denotes the sequence of observed outcomes and *a*_1:*T*_ the selected actions. *N* = 30 neural SSMs were trained independently for 50000 episodes. Optimization was performed using stochastic gradient descent (RMS-prop optimizer), minimizing the negative log-likelihood.

### Optimal Bayesian inference in the two-armed bandit task

For comparison, we implemented a normative Bayesian model that performs joint inference over the latent state and the unknown generative parameters of the environment (volatility and emission distributions). The optimal model assumes knowledge of the generative structure but not of the true parameter values. It performs Bayesian inference over: 1. the latent state *z*_*t*_, 2. the state volatility *h*, and 3. the mean *µ* and standard deviation *σ* parameters of the emission distribution.

Inference over the latent state *z*_*t*_was implemented using likelihood recursion^23^, allowing to computation of the posterior *p*(*z*_*t+1*_ | *y*_1:*t*_, *h, µ, σ*). Inference over the 3 parameters *h, µ* and *σ* was performed using a particle filtering approach^15,67^, with *N* = 100 particles. Increasing the number of particles to *N* = 1000 did not change performance at the fourth decimal place, indicating convergence of the approximation. Based on this, we used *N* = 100 particles for all optimal-model simulations across tasks.

Decisions in the optimal model were based on a particle filter approximation of the posterior distribution over the latent state, obtained by marginalizing over parameters:

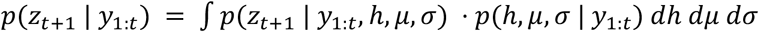

### Stimulus prediction task

On each trial *t*, a stimulus *z*_*t*_∈ {red, green} was sampled uniformly:

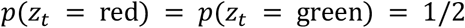

Conditional on the sampled stimulus, a stream of *N* symbols (cues) was generated independently. Each symbol in the stream was drawn from a stimulus-dependent distribution over four possible symbols.

For each symbol *k* ∈ {1, 2, 3, 4}, the probability that it was generated under the red stimulus was defined by a mapping probability *p*_*k*_ ∈ {0.2, 0.4, 0.6, 0.8}. Thus, the stimulus-dependent distributions were defined as:

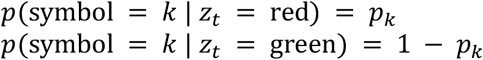

Given stimulus identity, each symbol of the observed symbol stream was therefore drawn from a multinomial distribution conditioned on *z*_*t*_.

After observing the symbol stream, the agent reported which stimulus had generated the stream. The true stimulus identity was then revealed. In contrast to the two-armed bandit task, the agent response did not influence the stimulus outcome.

Cue-stimulus associations (i.e., the mapping probabilities *p*_*k*_) could change over time. At each trial, mappings were resampled with probability *ρ*, where *ρ* was uniformly drawn from the set {0, 0.01, 0.03, 0.06, 0.1}. These changes required sustained learning and updating of the associative structure.

### Neural SSM architecture in the stimulus prediction task

The model maintained a latent state *z*_*t*_∈ {red, green} encoding stimulus identity on trial t. Inference over the latent state was performed by accumulating log-odds evidence across the observed symbol stream *C*_*t*_. Given a stream of observed cues *C*_*t*_, the posterior belief over stimulus identity was computed as:

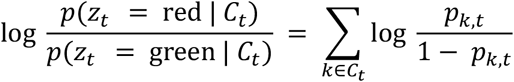

where *p*_*k,t*_ denotes the stimulus-specific conditional probability, *p*_*k,t*_ = *p*_ψ_ (symbol = k |z_*t*_ = red), with *ψ* the parameters of an associative network. The probabilities *p*_*k,t*_ were thus parameterized by a recurrent associative network implemented as a GRU network with 32 hidden units. At each trial, the network received as input: 1. the stimulus identity from the previous trial, and 2. a vector of dimension *K* = 4 encoding the counts of each symbol (cue) in the previous symbol stream.

This input allowed the network to update its internal representation of cue–stimulus associations based on trial history. The GRU hidden state was mapped to a four-dimensional output vector and passed through a sigmoid nonlinearity to obtain 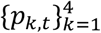, representing the current estimates of *p*_*k,t*_= *P*_ψ_ (symbol = *k*|*z*_*t*_= red) for each symbol.

These learned probabilities were then used in the log-odds accumulation step to compute the posterior belief over stimulus identity.

### Neural SSM training in the stimulus prediction task

Model parameters *ψ* were optimized by maximizing the joint log-likelihood of true stimulus identities and the symbol streams:

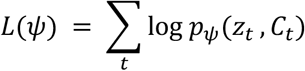

*N* = 30 neural SSMs were trained independently for 50,000 episodes. Optimization was performed using stochastic gradient descent (RMS-prop optimizer), minimizing the negative log-likelihood.

### Optimal Bayesian inference in the stimulus prediction task

The optimal Bayesian model assumes knowledge of the generative structure and of the possible values of the symbol–stimulus associations *p*_*k*_ ∈ {0.2, 0.4, 0.6, 0.8}, but not of the volatility parameter *ρ* governing changes in the associations. The goal of the agent is to infer the latent stimulus *z*_*t*_ on each trial from the observed symbol stream *C*_*t*_.

To do so, the optimal model introduces a latent variable *q*_*t*_ that encodes the current symbol-stimulus mapping, assigning to each symbol *k* a probability *p*_*k*_ ∈ {0.2, 0.4, 0.6, 0.8}. Inference over *q*_*t*_ was implemented using the likelihood recursion, allowing computation of the posterior *p*(*q*_*t*+1_ | *z*_1:*t*_, *C*_1:*t*_, *ρ*) with *C*_1:*t*_ the previously observed symbol streams.

Inference over the volatility parameter *ρ*, which governs changes *q*_*t*_, was performed using a particle filtering approach.

Regarding decision, given a new symbol stream *C*_*t+1*_, the model computes the posterior over the stimulus by marginalizing over the latent mappings:

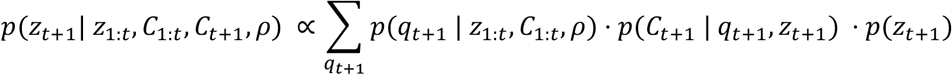

where *p*(*z*_*t+1*_) = 1/2. The likelihood term *p*(*C*_*t+1*_ | *q*_*t+1*_, *z*_*t+1*_) is computed by combining the symbol-wise probabilities under the multinomial observation model (implemented in log space). Finally, the posterior *p*(*z*_*t+1*_| *z*_*t+t*_, *C*_*t+t+1*_) over the stimulus is obtained by marginalizing over the volatility parameter – as in the optimal Bayesian model of the two-armed bandit task.

### Mastermind task

To evaluate the performance of neural SSMs in a task environment with richer combinatorial structure, we considered a variant of the Mastermind task. In this task, the environment contains a latent combination between a set of stimuli and a set of actions. At each trial *t*, the agent observes a stimulus *x* _*t*_drawn uniformly from a set of *N* = 4 stimuli. The correct action 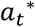 is determined by a latent permutation mapping between stimuli and actions. The set of possible combinations therefore corresponds to the set of permutations of the four actions, resulting in *K* = *N*! = 4!= 24 possible latent states.

The latent combination evolves over time according to a Markov process. At each trial, the current combination persists with probability 1 − *h* and switches to a different permutation with probability *h*, where *h* is the volatility parameter governing the rate of environmental change. When a switch occurs, a new combination is sampled uniformly among all permutations excluding the previous one.

After observing stimulus *x*_*t*_, the agent selects the action *a*_*t*_. Feedback *y*_*t*_is provided indicating whether the chosen action matches the correct action 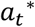 under the current combination. However, feedback is noisy: with probability 1−*f*, the feedback correctly indicates the corresponding outcome (true positive or true negative), whereas with probability *f*, the misleading outcome is shown (false positive or false negative). The parameter *f* therefore controls emission uncertainty in the task.

For each simulated environment, the volatility parameter *h* and the false-feedback rate *f* were sampled from truncated exponential distributions and remained fixed within a task instance.

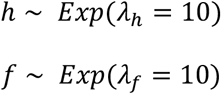

with *h* truncated between [0, 0.2] and *f* truncated between [0, 0.4].

### Neural SSM architecture in the Mastermind task

The neural state-space model used in the Mastermind task followed the architecture described for the two-armed bandit task, with modifications to accommodate the larger latent state space. In this setting, the latent state *z*_*t*_ encoded the current stimulus–action combination, corresponding to one of the *K* = 4!= 24 possible permutations between the four stimuli and the four actions. As in the reversal-learning task, the model maintained the joint likelihood *p*(*z*_*t*_, *y*_1:*t*_| *x*_1:*t*_, *a*_1:*t*_) and updated the posterior belief *p*(*z*_*t*_| *y*_1:*t*_, *x*_1:*t*_, *a*_1:*t*_) using likelihood recursion^23^.

After observing stimulus *x*_*t*_, the agent selected the action corresponding to the most probable correct response under its current belief over combinations:

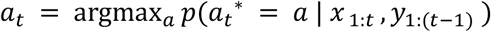

where the probability of the correct action is obtained by marginalizing over the posterior distribution over latent combinations.

Emission uncertainty, reflecting the probability of receiving correct or incorrect feedback, was modeled by a recurrent emission network *p*_θ_. The emission module was also implemented as a GRU network with 32 hidden units and produced a Bernoulli parameter governing the probability that feedback correctly indicated whether the chosen action matched the combination. The emission network received as input the current filtering belief of the selected combination and the observed reward signal, allowing it to adaptively estimate the reliability of feedback from experience.

Transition uncertainty over the latent combination was captured by a recurrent transition network that estimated the switching probability *h*_*t*_ from trial history. The transition module was implemented as a GRU network with 32 hidden units, whose hidden state was mapped to a scalar volatility parameter through a linear projection followed by a sigmoid nonlinearity. After observing outcome *y*_*t*_following action *a*_*t*_, the network received as input the log-probability that of the observed outcome if the chosen action matched the best action: log *p*_θ_ (*y*_*t*_). This parameter determined the probability that the latent combination switched to a different permutation at the next trial.

### Optimal Bayesian inference in the Mastermind task

The optimal Bayesian model assumes knowledge of the generative structure but not of the true parameter values. It performs Bayesian inference over: 1. the latent state *z*_*t*_, which encodes the current stimulus–action combination, corresponding to one of the *K* = 4! = 24 possible permutations between the four stimuli and the four actions, 2. the state volatility *h*, and 3. the false feedback rate *f*.

Inference over the latent state *z*_*t*_was implemented using likelihood recursion^23^, allowing computation of the posterior *p*(*z*_*t+1*_ | *x*_1:*t*_, *y*_1:*t*_, *h, f*), where *x*_1:*t*_ and *y*_1:*t*_ denote the previous observed stimuli and feedback. Inference over the parameters *h* and *f* was performed using a particle filtering approach^15,67^.

As for the two-armed bandit task, decisions in the optimal Bayesian model were based on the posterior distribution over the latent state *p*(*z*_*t*+1_ | *y*_1:*t*_, *x*_1:*t*_), obtained by marginalizing over parameters *h* and *f*. For a given stimulus *x*_*t*+1_, the posterior probability of each action was obtained by summing the posterior mass of all combinations *z*_*t*+1_ that assign that stimulus to that action. The optimal Bayesian model then selected the action with the highest posterior probability. Additional details are provided in a published study^10^.

### Category prediction task

The category prediction task can be viewed as a variant of the stimulus prediction task in which feedback about the true stimulus is noisy. On each trial t, a latent category *z*_*t*_ ∈ {red, green} was sampled uniformly and was never directly revealed to the agent. The category generated two types of observations.

First, as in the stimulus prediction task, a sequence of *N* discrete symbols was drawn according to category-specific distributions defined by symbol–category associations. These associations were unknown to the agent and could change over time with a fixed hazard rate, requiring sustained learning of the associative structure.

Second, after the agent made a category prediction based on the symbol stream, a continuous stimulus was presented as noisy feedback about the true category. The stimulus lay on a one-dimensional axis between red and green and was sampled from overlapping category-specific distributions. The degree of overlap was controlled by an emission-uncertainty parameter *f*, such that larger values produced more ambiguous feedback. Because the true category was never revealed, learning relied solely on probabilistic feedback from this noisy stimulus.

Performance was evaluated as the proportion of trials on which the category predicted by the agent matched the true latent category.

### Neural SSM architecture in the category prediction task

The neural SSM architecture re-used previously learned components from earlier tasks without fine-tuning. Specifically, we reused: 1. the associative component trained in the stimulus prediction task and 2. the emission component trained in the two-armed bandit task. Because *N* = 30 neural SSMs were trained independently in each of those tasks, we constructed N = 30 neural SSMs for the category prediction task by pairing one associative module and one emission module drawn from the corresponding trained agents.

In the original stimulus prediction task, the associative component received as input: 1. the stimulus identity from the previous trial, and 2. a vector of dimension *N*_symbols_ encoding the counts of each symbol in the previous symbol stream.

In the category prediction task, the true stimulus identity is never observed. Therefore, in the emission-reuse configuration, the stimulus input was replaced by an evidence signal derived from the emission component. Specifically, given observed noisy feedback *y*_*t*_, we provided the log-likelihood ratio between the two category hypotheses:

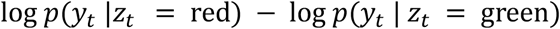

This quantity was transformed using a hyperbolic tangent nonlinearity to produce a bounded signal in [−1,1], which served as a proxy for feedback about category identity. Together with the symbol-count vector, this input allowed the associative network to update its internal representation of symbol–category associations. Crucially, in the reuse configuration, the association network was not retrained or adapted to this proxy signal; it was used exactly as trained in the stimulus prediction task.

The emission component provided probabilistic information about the relationship between latent category and observed continuous stimulus. At each trial, it received as input: 1. the current posterior belief over the selected category from the previous trial *p*(*z*_*t*_= *a*_*t*_| *y*_*t+*(*t*-1)_) – where the chosen category was *a*_*t*_= 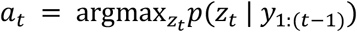 – and 2. the observed noisy stimulus *y*_*t*_.

### Optimal Bayesian inference in the category prediction task

The optimal Bayesian model assumes knowledge of the generative structure and of the possible values of the symbol–category associations *p*_*k*_ ∈ {0.2, 0.4, 0.6, 0.8}, but not of the volatility parameter *ρ* governing changes in the associations, nor of the mean *µ* and the standard deviation *σ* of the category-specific feedback distribution. The goal of the agent is to infer the latent category *z*_*t*_on each trial from the observed symbol stream *C*_*t*_.

As in the stimulus prediction task, the optimal Bayesian model introduces a latent variable *q*_*t*_ that encodes the current symbol-category mapping, assigning to each symbol *k* a probability *p*_*k*_ ∈ {0.2, 0.4, 0.6, 0.8}. Inference over *q*_*t*_was implemented using the likelihood recursion, allowing computation of the posterior *p*(*q*_*t+1*_ | *y*_1:*t*_, *C*_1:*t*_, *ρ, µ, σ*) with *C*_1:*t*_ and *y*_1:*t*_ the previously observed symbol streams and feedbacks.

Inference over the volatility parameter *ρ, µ* and *σ* was performed using a particle filtering approach. Regarding decision, given a new symbol stream *C*_*t+1*_, the model computes the posterior over the category by marginalizing over the latent mappings:

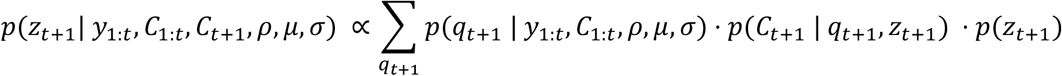

where *p*(*z*_*t+1*_) = 1/2. The likelihood term *p*(*C*_*t+1*_ | *q*_*t+1*_, *z*_*t+1*_) is computed by combining the symbol-wise probabilities under the multinomial observation model (implemented in log space). Finally, the posterior *p*(*z*_*t+1*_| *y*_1:*t*_, *C*_1:*t+1*_) over the category, used for decision, is obtained by marginalizing over the latent parameters.

### History-dependent category prediction task

The history-dependent category prediction task extends the category prediction task by introducing temporal dependencies in the latent category. Instead of being sampled independently across trials, the category evolves according to a stochastic process, creating transition uncertainty in addition to associative and emission uncertainty.

On each trial *t*, a latent category *z*_*t*_ ∈ {red, green} governed the observations but was never directly revealed to the agent. The category evolved across trials according to a two-state Markov process with volatility parameter *h*:

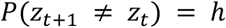

so that larger values of *h* produced more frequent category switches. This introduced history dependence, requiring agents to track the current category over time.

### Neural SSM architecture in the history-dependent category prediction task

The neural state-space model reused recurrent components previously trained in earlier tasks. The model combined three components (GRU networks): 1. a transition network estimating category volatility, 2. an emission network modeling the mapping from latent category to noisy stimulus feedback, and 3. an association network encoding symbol–category relationships.

### Optimal Bayesian inference in the history-dependent category prediction task

The optimal model extends the category prediction model by incorporating temporal dependencies in the latent category. In addition to the volatility parameter *ρ* governing changes in symbol–category associations and the emission parameters *µ* and *σ*, the model infers a transition volatility parameter *h* controlling the Markovian dynamics of the latent category.

Inference over the latent variables *z*_*t*_(category) and *q*_*t*_(symbol–category mapping) was implemented using likelihood recursion, applied over the joint state space of category and mapping, allowing computation of the posterior *p*(*z*_*t+1*_, *q*_*t+1*_ | *y*_1:*t*_, *C*_1:*t*_, *h, ρ, µ, σ*). Inference over the parameters *h, ρ, µ* and *σ* was performed using a particle filtering approach.

Regarding decision, given a new symbol stream *C*_*t+1*_, the model computes the posterior over the category by combining prior beliefs induced by the transition dynamics with evidence from the symbol stream, and marginalizing over the latent mappings. The posterior over the category is then obtained by marginalizing over the latent parameters.

### Structure matching between the bandit and apples tasks

The bandit and apples tasks tested in human participants were designed to have the same latent structure as the two-armed bandit task described above. The bandit task corresponded to the previously described two-armed bandit task, where a latent binary state *z*_*t*_ ∈ {0, 1} encoded which symbol was currently the higher-rewarding option – associated with a probability distribution of rewards between 1 and 99 points. The apples task corresponded to a sequential two-alternative forced-choice (2-AFC) visual categorization task where the latent binary state *z*_*t*_ ∈ {0, 1} encoded which source – associated with a probability distribution of colors along a red-green hue continuum – currently generated the presented apples.

The two tasks were perfectly matched by using the same probability distributions for sampling rewards in the bandit task, and color hues in the apples task, and by making the rewards associated with the two options sum to 100 on every single trial, as in the two-armed bandit task described above. The false-feedback rate *f* was set to 0.30 in the bandit task, and the category overlap *f* was therefore set to the same value (0.30) in the apples task.

Reversals of the latent binary state in the two tasks occurred with a fixed state volatility *h* = 0.11, and controlled by episodes whose length (in trials) was sampled from a truncated exponential distribution of mean 12 trials, minimum 6 trials and maximum 24 trials. A reversal in the bandit task triggered a change in the symbol corresponding to the better (higher-rewarding) option, whereas a reversal in the apples task triggered a change in the source generating the presented apples.

### Human experimental conditions

Adult neurotypical participants with no history of neurological nor psychiatric conditions were recruited online on the Prolific.co platform. Upon gaining access to the study, participants were prompted to set the task to full-screen before being able to continue. Next, participants were introduced to the mechanics of the two tasks. They were informed that they would take part in two separate games (the bandit and apples tasks) divided into 4 runs of 72 trials each (total: 288 trials per task), without emphasizing the similarities or differences between them. In a counterbalanced fashion, participants performed training in one of the two tasks – either the bandit or apples task – before performing a practice run of this first task. Then, they performed training on the other task before playing a practice run of this second task. The training runs familiarized participants with the specifics of each task, and instructed them on the payout scheme associated with each task. Participants then alternated between runs of the two tasks (1-A, 2-B, 3-A, 4-B, 5-A, 6-B, 7-A, 8-B), with short self-paced breaks between successive runs. Participants replayed the exact same experiment with the same instances of the two tasks in the retest session, two weeks (± 1 day) later.

At the end of the experiment, participants completed the 16-item International Cognitive Ability Resource Test (ICAR-16). Total completion of the study took approximately 50 minutes. Participants were compensated £6.25 for completion of the test session, £7.50 for completion of the retest session, and could obtain bonuses of £1.00 or £2.00 depending on their overall performance – the fraction of trials in which they chose the higher-rewarding option in the bandit task, or the correct source generating the presented apples in the apples task.

All participants provided informed consent regarding their participation in the study, which followed guidelines approved by the ethical review committee of the Institut National de la Santé et de la Recherche Médicale (IRB #00003888). We excluded from analyses any participant whose accuracy did not significantly exceed chance-level performance at a one-tailed threshold α = 0.05 in any of the two tasks (binomial test against chance). In total, this exclusion criterion retained 129 participants (mean age of 30 ± 6 years) who completed the test and retest sessions.

### Human behavioral modeling

We modelled human behavior in the bandit and apples task by a previously validated hidden-state inference model that tracks the latent binary state *z*_*t*_of each task from observations *x*_1:*t*_ – rewards in the bandit task, colored apples in the apples task^29–32^. The model relies on the normative equation to update its probabilistic belief *L*_*t*_ regarding the current latent binary state:

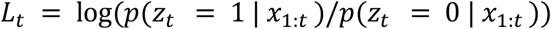

The probabilistic belief *L*_*t*_accounts for a subjective, fixed state volatility *h*:

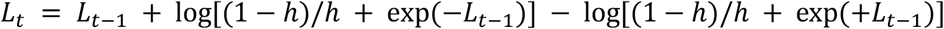

The probabilistic belief *L*_*t*_ is then updated using the evidence *y*_*t*_ provided by the new observation *x*_*t*_ – the obtained reward in the bandit task, the color of the presented apple in the apples task:

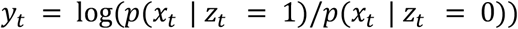

yielding *L*_*t*_→ *L*_*t*_+ *y*_*t*_.

Following known biases in reward-guided and perceptual decision-making, we modeled possible distortions in the evidence function *F*(·) which maps observations *x*_*t*_ onto evidence *y*_*t*_using additional, task-specific parameters. In the bandit task, we modeled loss aversion using a reference point parameter *r*_F_ and a loss aversion strength *α* following prospect theory^33,34^:

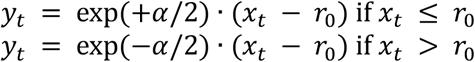

with *x*_*t*_ the observed reward between 1 and 99 points, and the distorted evidence *y*_*t*_rescaled in the same range as the non-distorted evidence. In the apples task, we modeled categorization distortions^35,36^ using a category bias *ω* and a category attraction strength *γ*:

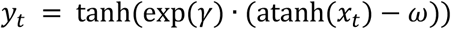

with *x*_*t*_ the color hue of the presented apple in [-1,+1] (from -1 for pure red to +1 for pure green), and the distorted evidence *y*_*t*_ rescaled in the same range as the non-distorted evidence.

The model chooses which action to take using a softmax policy appropriate for each task: selecting the higher-rewarding option given the most likely latent state *z*_*t*_in the bandit task (*z*_*t*_= 1 if *L*_*t*_> 0, *z*_*t*_= 0 if *L*_*t*_< 0), and selecting the current source drawing the apples given the most likely latent state *z*_*t*_ in the apples task. This softmax policy is controlled by a temperature parameter τ.

We fitted this four-parameter model – {*h, α, r*_0_, τ} in the bandit task, {*h, γ, ω*, τ} in the apples task – to each participant’s choices made in the two tasks. The log-likelihoods associated with each action were estimated using a weighted bootstrap-based particle filter, and then combined with prior distributions on parameter values to get an unnormalized log-posterior estimate. The prior distributions over parameter values were set as follows: uniform between 0 and 1 for *h* (same across tasks), exponential of mean 1 for τ (same across tasks), normal of mean 0 and s.d. 1 for *α* (bandit task) and *γ* (apples task), and normal of mean 0 and s.d. 0.1 for *r*_0_ (bandit task) and *ω* (apples task). This unnormalized log-posterior was given as argument to the Variational Bayesian Monte Carlo (VBMC) algorithm^68^ returning a variational approximation of the full posterior and a lower bound on the log-marginal likelihood. We took the posterior mean (and standard deviation) to obtain best-fitting parameter values (and standard deviation).

We verified the ability of the model to provide accurate estimates of each parameter through parameter recovery by correlating each best-fitting parameter to the parameter values used for simulation. Recovered parameter values for *h* and *τ* were found to be highly correlated to ground-truth parameters in both tasks (**Supplementary Fig. 7**).

## Acknowledgments

This research was funded by a grant from the Agence Nationale de la Recherche awarded to V.W. (MONODEC, ANR-23-CE37-0028), an institutional grant from the Agence Nationale de la Recherche awarded to the Département d’Études Cognitives (FrontCog, ANR-17-EURE-0017), a grant from the Swiss National Science Foundation awarded to A.P. (10.004.051), and a Royal Society Newton International Fellowship awarded to J.B. (NIF\R1\252778). We thank Felix Hubert and Peter Latham for useful discussions.

## Code and data availability

Code for the neural SSMs is publicly available at https://github.com/csmfindling/neuralSSMs. Human behavioral data and analysis code (including model fitting) for the bandit and apples tasks are publicly available at https://doi.org/10.5281/zenodo.20025309. Human behavioral for the stable and volatile bandit conditions are publicly available at https://doi.org/10.5281/zenodo.7624899.

## Contributions

Study conception: C.F., V.W. Experiment design and data acquisition: C.F., J.K.L., V.W. Data analysis and interpretation: C.F., V.W. Manuscript writing: C.F., J.J.W.B., A.P., V.W.

## References

1. Knill, D. C. & Pouget, A. The Bayesian brain: the role of uncertainty in neural coding and computation. Trends Neurosci. 27, 712–719 (2004).

2. Körding, K. P. & Wolpert, D. M. Bayesian integration in sensorimotor learning. Nature 427, 244–247 (2004).

3. Ma, W. J., Beck, J. M., Latham, P. E. & Pouget, A. Bayesian inference with probabilistic population codes. Nat Neurosci 9, 1432–1438 (2006).

4. Rao, R. P. & Ballard, D. H. Predictive coding in the visual cortex: a functional interpretation of some extra-classical receptive-field effects. Nat. Neurosci. 2, 79–87 (1999).

5. Friston, K. A theory of cortical responses. Philos. Trans. R. Soc. Lond., B, Biol. Sci. 360, 815–836 (2005).

6. Bayesian Brain: Probabilistic Approaches to Neural Coding. (The MIT Press, 2006). doi:10.7551/mitpress/9780262042383.001.0001.

7. Perception as Bayesian Inference. (Cambridge University Press, Cambridge, 1996). doi:10.1017/CBO9780511984037.

8. Oaksford, M. & Chater, N. Bayesian Rationality: The Probabilistic Approach to Human Reasoning. (Oxford University Press, 2007).

9. Tenenbaum, J. B., Griffiths, T. L. & Kemp, C. Theory-based Bayesian models of inductive learning and reasoning. Trends Cogn Sci 10, 309–318 (2006).

10. Findling, C., Chopin, N. & Koechlin, E. Imprecise neural computations as a source of adaptive behaviour in volatile environments. Nat Hum Behav 5, 99–112 (2021).

11. Piray, P. & Daw, N. D. Computational processes of simultaneous learning of stochasticity and volatility in humans. Nat Commun 15, 9073 (2024).

12. Findling, C., Romand-Monnier, M., Skvortsova, V. & Koechlin, E. Neural variability in the medial prefrontal cortex contributes to efficient adaptive behavior. Nat Commun 16, 11356 (2025).

13. Beal, M. J. Variational algorithms for approximate Bayesian inference. (UCL (University College London), 2003).

14. Andrieu, C., Doucet, A. & Holenstein, R. Particle Markov Chain Monte Carlo Methods. Journal of the Royal Statistical Society Series B: Statistical Methodology 72, 269–342 (2010).

15. Chopin, N., Jacob, P. E. & Papaspiliopoulos, O. SMC 2 : an efficient algorithm for sequential analysis of state space models. Journal of the Royal Statistical Society. Series B (Statistical Methodology) 75, 397–426 (2013).

16. Gershman, S. & Goodman, N. Amortized Inference in Probabilistic Reasoning. Proceedings of the Annual Meeting of the Cognitive Science Society 36, (2014).

17. Kingma, D. P. & Welling, M. Auto-Encoding Variational Bayes. Preprint at https://doi.org/10.48550/arXiv.1312.6114 (2022).

18. Heald, J. B., Lengyel, M. & Wolpert, D. M. Contextual inference underlies the learning of sensorimotor repertoires. Nature 600, 489–493 (2021).

19. Yang, G. R., Joglekar, M. R., Song, H. F., Newsome, W. T. & Wang, X.-J. Task representations in neural networks trained to perform many cognitive tasks. Nat. Neurosci. 22, 297–306 (2019).

20. Driscoll, L. N., Shenoy, K. & Sussillo, D. Flexible multitask computation in recurrent networks utilizes shared dynamical motifs. Nat Neurosci 27, 1349–1363 (2024).

21. Riveland, R. & Pouget, A. Natural language instructions induce compositional generalization in networks of neurons. Nat Neurosci 27, 988–999 (2024).

22. Bakermans, J. J. W., Tano, P., Riveland, R., Findling, C. & Pouget, A. Compositional meta-learning through probabilistic task inference. Preprint at 10.48550/arXiv.2510.01858 (2025).

23. Scott, S. L. Bayesian Methods for Hidden Markov Models: Recursive Computing in the 21st Century. Journal of the American Statistical Association 97, 337–351 (2002).

24. Knowlton, B. J., Mangels, J. A. & Squire, L. R. A neostriatal habit learning system in humans. Science 273, 1399–1402 (1996).

25. Poldrack, R. A. et al. Interactive memory systems in the human brain. Nature 414, 546–550 (2001).

26. Gluck, M. A., Shohamy, D. & Myers, C. How do people solve the ‘weather prediction’ task?: individual variability in strategies for probabilistic category learning. Learn. Mem. 9, 408–418 (2002).

27. Yang, T. & Shadlen, M. N. Probabilistic reasoning by neurons. Nature 447, 1075–1080 (2007).

28. Donoso, M., Collins, A. G. E. & Koechlin, E. Foundations of human reasoning in the prefrontal cortex. Science 344, 1481–1486 (2014).

29. Glaze, C. M., Kable, J. W. & Gold, J. I. Normative evidence accumulation in unpredictable environments. eLife 4, 08825 (2015).

30. Glaze, C. M., Filipowicz, A. L. S., Kable, J. W., Balasubramanian, V. & Gold, J. I. A bias– variance trade-off governs individual differences in on-line learning in an unpredictable environment. Nat. Hum. Behav. 2, 213–224 (2018).

31. Weiss, A., Chambon, V., Lee, J. K., Drugowitsch, J. & Wyart, V. Interacting with volatile environments stabilizes hidden-state inference and its brain signatures. Nat. Commun. 12, 2228 (2021).

32. Drevet, J., Drugowitsch, J. & Wyart, V. Efficient stabilization of imprecise statistical inference through conditional belief updating. Nat Hum Behav 6, 1691–1704 (2022).

33. Kahneman, D. & Tversky, A. Prospect Theory: An Analysis of Decision under Risk. Econometrica 47, 263–291 (1979).

34. Tversky, A. & Kahneman, D. Loss Aversion in Riskless Choice: A Reference-Dependent Model. The Quarterly Journal of Economics 106, 1039–1061 (1991).

35. Huttenlocher, J., Hedges, L. V. & Duncan, S. Categories and particulars: prototype effects in estimating spatial location. Psychol Rev 98, 352–376 (1991).

36. Goldstone, R. L. Effects of Categorization on Color Perception. Psychological Science 6, 298–304 (1995).

37. Wilson, R. C., Geana, A., White, J. M., Ludvig, E. A. & Cohen, J. D. Humans use directed and random exploration to solve the explore-exploit dilemma. J. Exp. Psychol. Gen. 143, 2074–2081 (2014).

38. Wilson, R. C., Bonawitz, E., Costa, V. D. & Ebitz, R. B. Balancing exploration and exploitation with information and randomization. Curr Opin Behav Sci 38, 49–56 (2021).

39. Gershman, S. J. Deconstructing the human algorithms for exploration. Cognition 173, 34–42 (2018).

40. Schulz, E. & Gershman, S. J. The algorithmic architecture of exploration in the human brain. Curr Opin Neurobiol 55, 7–14 (2019).

41. Lee, J. K., Rouault, M. & Wyart, V. Adaptive tuning of human learning and choice variability to unexpected uncertainty. Sci Adv 9, eadd0501 (2023).

42. Krishnan, R. G., Shalit, U. & Sontag, D. Structured Inference Networks for Nonlinear State Space Models. Preprint at 10.48550/arXiv.1609.09869 (2016).

43. Ashwood, Z. C. et al. Mice alternate between discrete strategies during perceptual decision-making. Nat Neurosci 25, 201–212 (2022).

44. Buesing, L., Bill, J., Nessler, B. & Maass, W. Neural Dynamics as Sampling: A Model for Stochastic Computation in Recurrent Networks of Spiking Neurons. PLoS Comput Biol 7, e1002211 (2011).

45. Berkes, P., Orbán, G., Lengyel, M. & Fiser, J. Spontaneous cortical activity reveals hallmarks of an optimal internal model of the environment. Science 331, 83–87 (2011).

46. Orbán, G., Berkes, P., Fiser, J. & Lengyel, M. Neural Variability and Sampling-Based Probabilistic Representations in the Visual Cortex. Neuron 92, 530–543 (2016).

47. Zhang, W.-H., Wu, S., Josić, K. & Doiron, B. Sampling-based Bayesian inference in recurrent circuits of stochastic spiking neurons. Nat Commun 14, 7074 (2023).

48. Wang, J. X. et al. Prefrontal cortex as a meta-reinforcement learning system. Nat. Neurosci. 21, 860–868 (2018).

49. Botvinick, M. et al. Reinforcement learning, fast and slow. Trends Cogn. Sci. 23, 408–422 (2019).

50. Vaswani, A. et al. Attention is all you need. in Advances in Neural Information Processing Systems (NeurIPS) 5998–6008 (2017).

51. Daw, N. D., O’Doherty, J. P., Dayan, P., Seymour, B. & Dolan, R. J. Cortical substrates for exploratory decisions in humans. Nature 441, 876–879 (2006).

52. Findling, C., Skvortsova, V., Dromnelle, R., Palminteri, S. & Wyart, V. Computational noise in reward-guided learning drives behavioral variability in volatile environments. Nat. Neurosci. 22, 2066–2077 (2019).

53. Usher, M. & McClelland, J. L. The time course of perceptual choice: the leaky, competing accumulator model. Psychol Rev 108, 550–592 (2001).

54. Ratcliff, R. & Smith, P. L. A comparison of sequential sampling models for two-choice reaction time. Psychol. Rev. 111, 333–367 (2004).

55. Bogacz, R., Brown, E., Moehlis, J., Holmes, P. & Cohen, J. D. The physics of optimal decision making: a formal analysis of models of performance in two-alternative forced-choice tasks. Psychol. Rev. 113, 700–765 (2006).

56. Moutoussis, M. et al. Decision-making ability, psychopathology, and brain connectivity. Neuron 109, 2025–2040.e7 (2021).

57. Schurr, R., Reznik, D., Hillman, H., Bhui, R. & Gershman, S. J. Dynamic computational phenotyping of human cognition. Nat Hum Behav 8, 917–931 (2024).

58. Zador, A. M. A critique of pure learning and what artificial neural networks can learn from animal brains. Nat Commun 10, 3770 (2019).

59. Lake, B. M., Ullman, T. D., Tenenbaum, J. B. & Gershman, S. J. Building machines that learn and think like people. Behav. Brain Sci. 40, e253 (2017).

60. Daw, N. D., Gershman, S. J., Seymour, B., Dayan, P. & Dolan, R. J. Model-based influences on humans’ choices and striatal prediction errors. Neuron 69, 1204–1215 (2011).

61. Starkweather, C. K., Babayan, B. M., Uchida, N. & Gershman, S. J. Dopamine reward prediction errors reflect hidden-state inference across time. Nat. Neurosci. 20, 581–589 (2017).

62. Babayan, B. M., Uchida, N. & Gershman, S. J. Belief state representation in the dopamine system. Nat Commun 9, 1891 (2018).

63. Blanco-Pozo, M., Akam, T. & Walton, M. E. Dopamine-independent effect of rewards on choices through hidden-state inference. Nat Neurosci 27, 286–297 (2024).

64. Behrens, T. E. J., Woolrich, M. W., Walton, M. E. & Rushworth, M. F. S. Learning the value of information in an uncertain world. Nat. Neurosci. 10, 1214–1221 (2007).

65. Constantinescu, A. O., O’Reilly, J. X. & Behrens, T. E. J. Organizing conceptual knowledge in humans with a gridlike code. Science 352, 1464–1468 (2016).

66. Mishchanchuk, K. et al. Hidden state inference requires abstract contextual representations in the ventral hippocampus. Science 386, 926–932 (2024).

67. Chopin, N. A Sequential Particle Filter Method for Static Models. Biometrika 89, 539–551 (2002).

68. Acerbi, L. Variational Bayesian Monte Carlo. in Advances in Neural Information Processing Systems vol. 31 (Curran Associates, Inc., 2018).

