## Supplementary Information for "Augmenting the Bayesian Brain with learned and reusable world-model components for flexible cognition"

### Contents

**Supplementary Figures 1-10** (pages 2-12)

**Supplementary Table 1** (page 13)

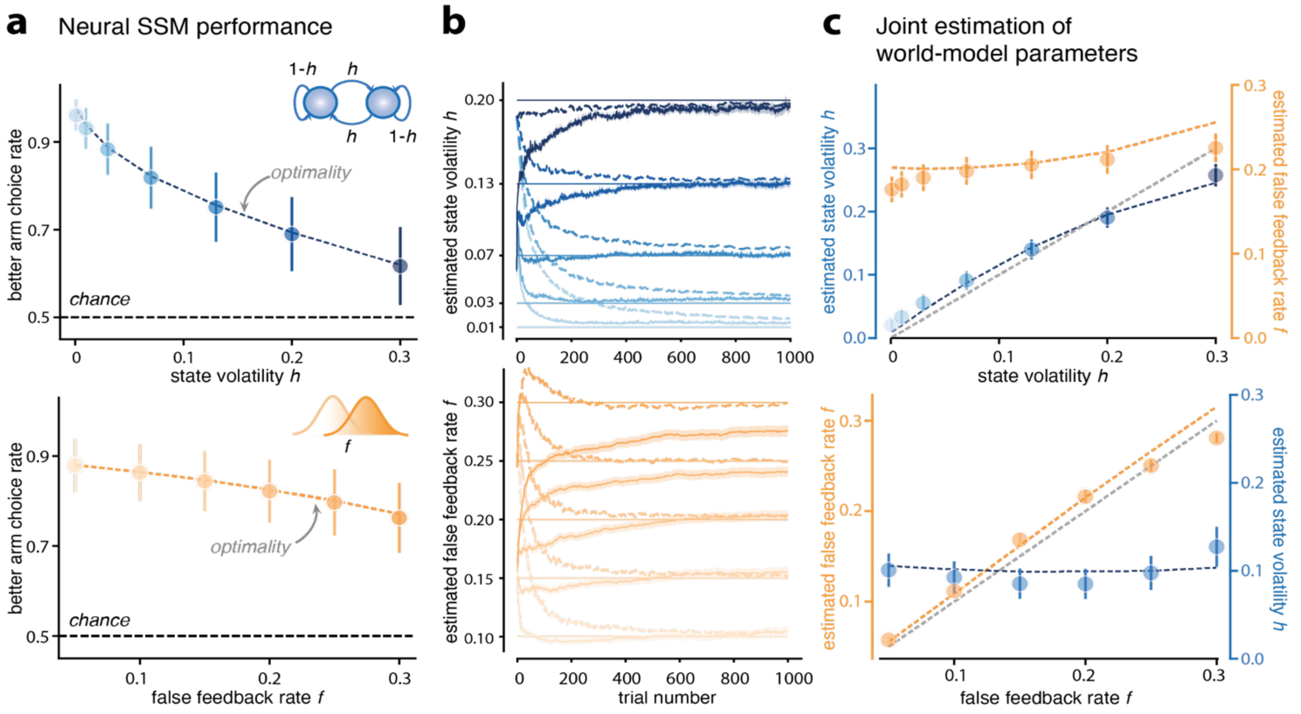


**Supplementary Figure 1 | Comparison of neural SSMs and the optimal Bayesian model in reversal learning** (**a**) Neural SSM performance relative to the optimal model. Top: Proportion of correct choices (better-arm choice rate) as a function of true state volatility *h*. Bottom: Proportion of correct choices as a function of true false-feedback rate *f*. In both cases, neural SSM performance closely matches the optimal Bayesian performance (dashed lines), showing that accuracy declines with increasing transition or emission uncertainty while remaining near-optimal. (**b**) Estimated world-model parameters. Top: Trial-by-trial estimates of state volatility for different true volatility levels (horizontal lines). Solid lines indicate neural SSM estimates, and dashed lines indicate estimates from the optimal Bayesian model. Bottom: Trial-by-trial estimates of the false-feedback rate for different true false-feedback levels (horizontal lines), with neural SSM estimates shown as solid lines and optimal-model estimates as dashed lines. (**c**) Joint estimation of world-model parameters. Top: Estimated state volatility and false-feedback rate as a function of true volatility. Bottom: Estimated false-feedback rate and state volatility as a function of true false-feedback rate. Colored dashed lines indicate parameter estimates obtained with the optimal Bayesian model. Neural SSM estimates closely match those of the optimal model, and transition and emission uncertainties remain dissociated, with each parameter varying selectively with its corresponding ground-truth value.

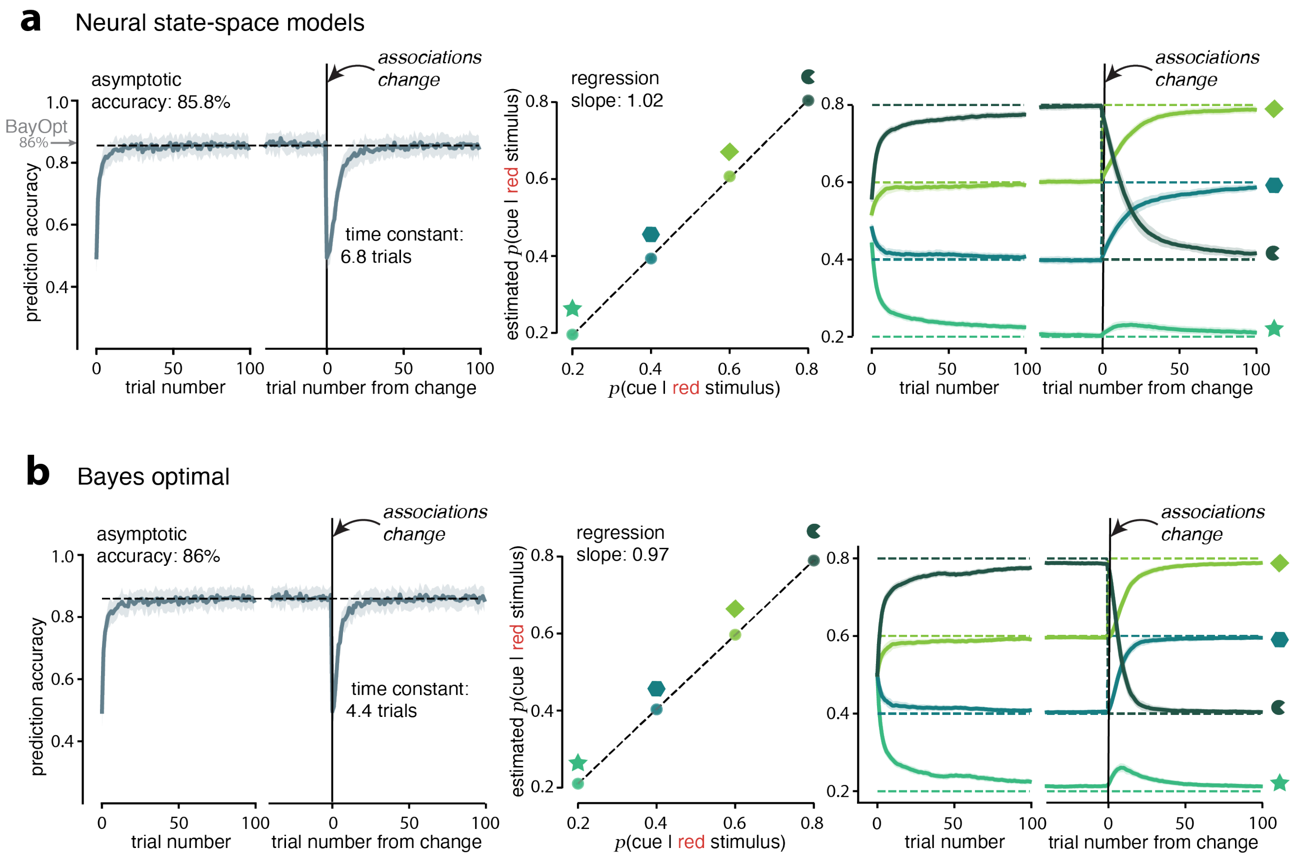


**Supplementary Figure 2 | Optimal Bayesian model in the stimulus prediction task** (**a**) Neural state-space models (reproduction of **Fig. 3ef**). Left: Stimulus prediction accuracy as a function of trial number and aligned to changes in symbol–stimulus associations. Middle: Relationship between true symbol–stimulus associations and the corresponding asymptotic amortized estimates of the neural SSM. Right: Time courses of learned associations, showing flexible updating of estimated probabilities before and after changes in symbol–stimulus mappings. (**b**) Bayes optimal observer. Same conventions as in (**a**). The optimal model exhibits comparable asymptotic accuracy (86%) and faster adaptation dynamics (time constant: 4.4 trials, compared to 6.8 trials for the neural-SSM), with accurate recovery of associative probabilities and flexible updating following associative changes.

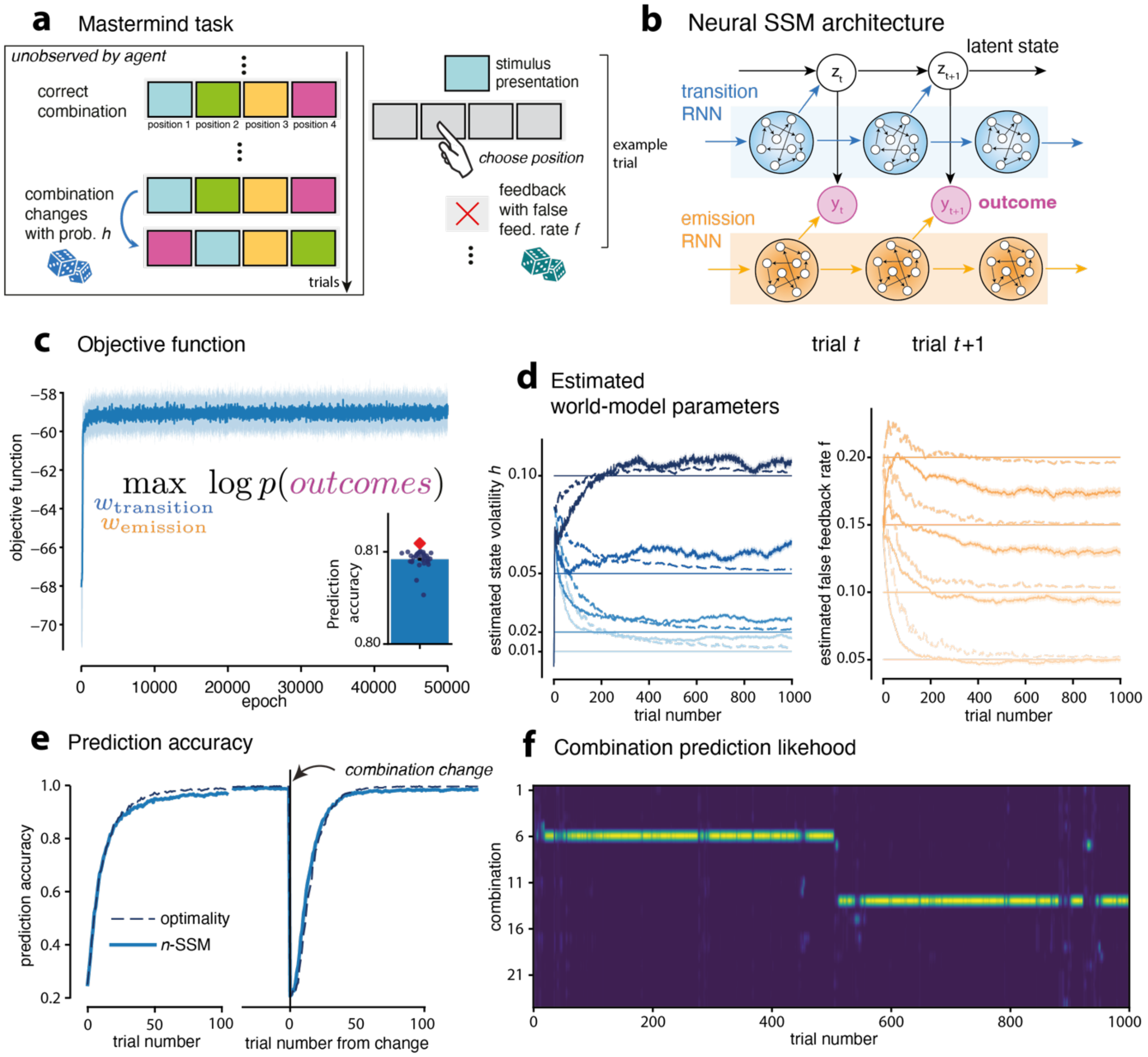


**Supplementary Figure 3 | Neural SSMs learn generative world models in the Mastermind task** (**a**) Mastermind task. On each trial, one of four stimuli is presented, and the agent must select the correct action according to a latent stimulus–action combination. The combination corresponds to one of the 24 possible permutations between stimuli and actions. After the agent's choice, binary feedback indicates whether the selected action matches the correct combination, with false feedback rate *f*. The latent combination may change over time according to a stochastic switching process, requiring sustained inference over the current combination. (**b**) Neural state-space model architecture. The neural SSM maintains a posterior distribution over the *24* possible stimulus–action combinations. As in the two-armed bandit task, the model combines two network modules; a transition network estimates the state volatility governing switches between combinations, while an emission network estimates the false-feedback rate. These quantities are used within the likelihood recursion to update beliefs over the latent combination. (**c**) Training dynamics. Log-likelihood of the observed feedback as a function of training episodes for *N=30* independently trained neural SSM agents. Training converges reliably across runs, indicating stable learning of the generative model. The bar plot summarizes final performance across agents, showing near-optimal accuracy (Bayes optimal performance shown in red). (**d**) Trial-by-trial estimates of the state volatility (left) and false-feedback rate (right) for different ground-truth parameter values (horizontal lines). Solid lines indicate neural SSM estimates, and dashed lines indicate estimates from the Bayes optimal model. (**e**) Choice accuracy aligned to switches in the latent combination. The neural SSM closely tracks the optimal adaptation dynamics (dashed line) following changes in the underlying stimulus–action combination, demonstrating sensitivity to transition uncertainty. (**f**) Posterior belief over combinations. Example posterior distribution over the 24 possible stimulus–action combinations across trials for a single task instance, with a combination switch at trial 500. The model quickly concentrates probability mass on the correct combination and shifts to a new combination following latent switches.

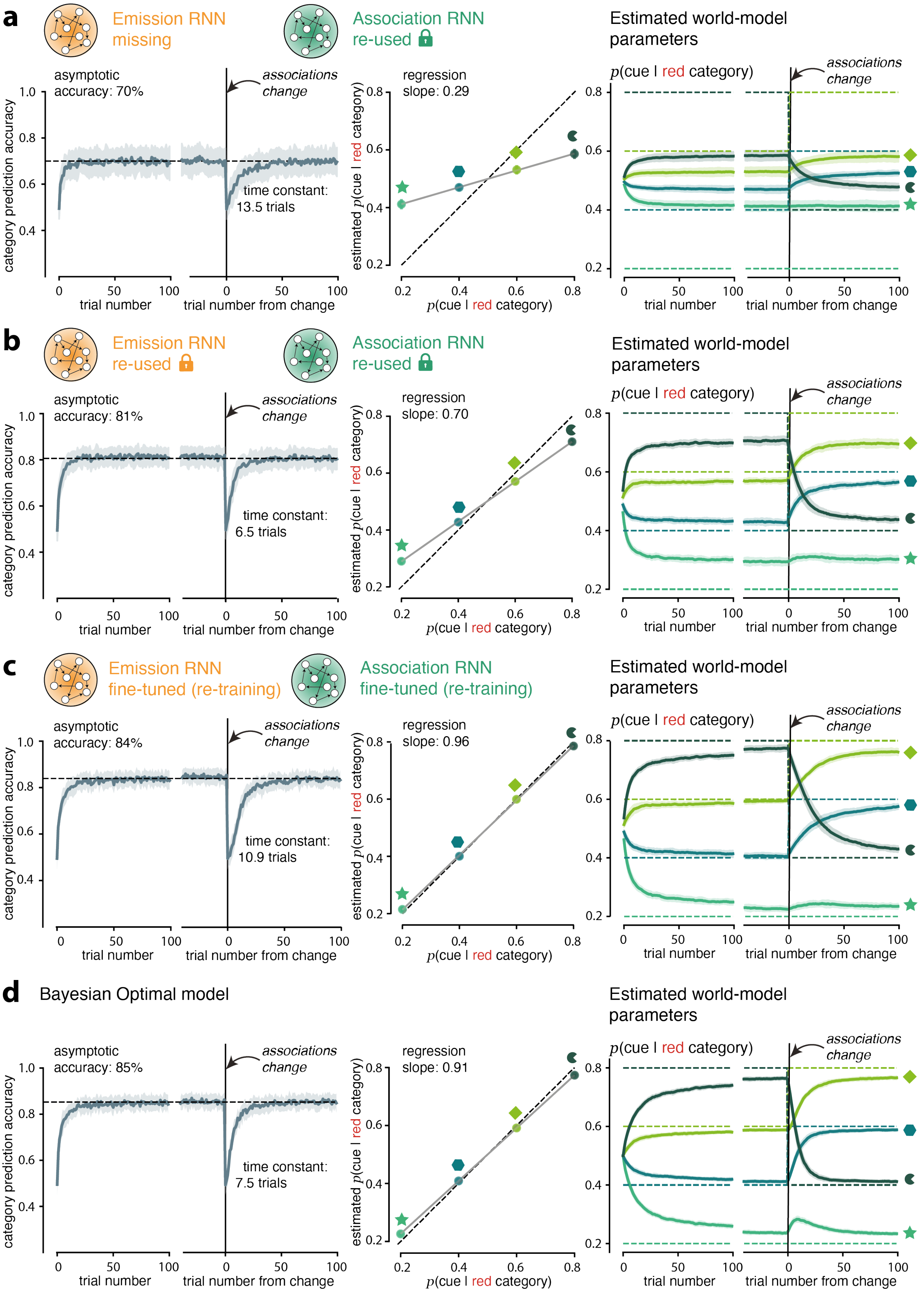


(figure caption on next page)

**Supplementary Figure 4 | Effect of world-model component reuse and fine-tuning in the category prediction task compared to the optimal Bayesian model** (**a**) reuse of the association network without an emission network, (**b**) reuse of both the emission and association networks without additional training, (**c**) reuse of the emission and association networks with additional fine-tuning and (**d**) the optimal Bayesian model. Left column: category prediction accuracy. Proportion of times that the latent category was correctly inferred as a function of trial number. Accuracy is shown during initial learning and aligned to changes in symbol–category associations, illustrating adaptation following associative changes. Middle column: recovery of associative probabilities. Relationship between true symbol–category associations and the corresponding asymptotic estimates produced by the association network. Each point corresponds to a symbol, demonstrating recovery of the underlying associative structure. Right column: trial-by-trial associative estimates. Time courses of stimulus-specific conditional probabilities $p(x_k|\, y = red)$ for each symbol. Vertical dashed lines indicate changes in the underlying symbol–category associations, and horizontal dashed lines indicate the true associative probabilities.

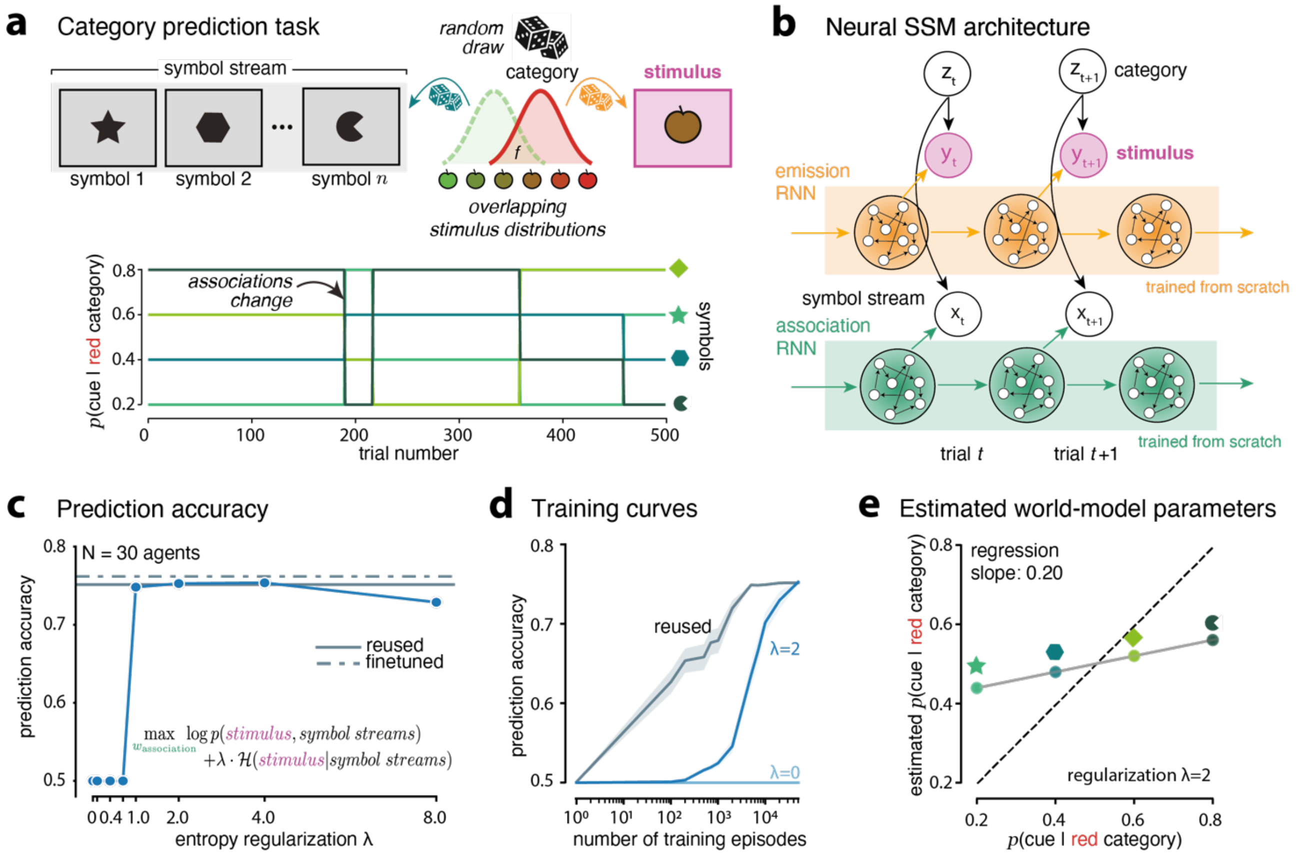


**Supplementary Figure 5 | Neural SSMs trained from scratch in the category prediction task** (**a**) Category prediction task, same as in Fig. 4 (**b**) Neural SSM architecture trained from scratch. The model consists of an emission network that estimates the distribution of the noisy stimulus given the category and an association network that learns symbol–category relationships. Both modules are trained jointly from scratch rather than reused from previous tasks. (**c**) Prediction accuracy after training. Category prediction accuracy following training for 50,000 episodes. An entropy regularization term was added to the loss function to encourage the association network to learn high-entropy solutions. Grey lines indicate the performance of the reused agent from Fig. 4, in which the emission and association networks are reused without fine-tuning. Dashed grey lines indicate the reused agent with additional fine-tuning of both modules. The reused agent performs virtually as well as the model trained from scratch with an optimized entropy regularization parameter. (**d**) Training curves. Prediction accuracy as a function of training episodes for models trained from scratch under different entropy regularization strengths ($\lambda$). Without regularization ($\lambda = 0$), the model remains at chance level. With stronger regularization ($\lambda = 2$), the model learns successfully. Grey curves show the performance of the reused agent constructed by combining partially trained modules: at training episode $T$, the emission network trained on the bandit task and the association network trained on the stimulus prediction task after $T$ episodes were stacked together to form the reused model. (**e**) Recovery of associative parameters. Relationship between true symbol–category associations and the estimates inferred by the association network. The slope of the regression (0.20) is substantially lower than that obtained for reused agents (Supplementary Fig. 4), indicating that models trained from scratch achieve good behavioral performance but recover the underlying world-model parameters less accurately; this arises because the entropy regularization encourages high-entropy solutions that support accurate category prediction.

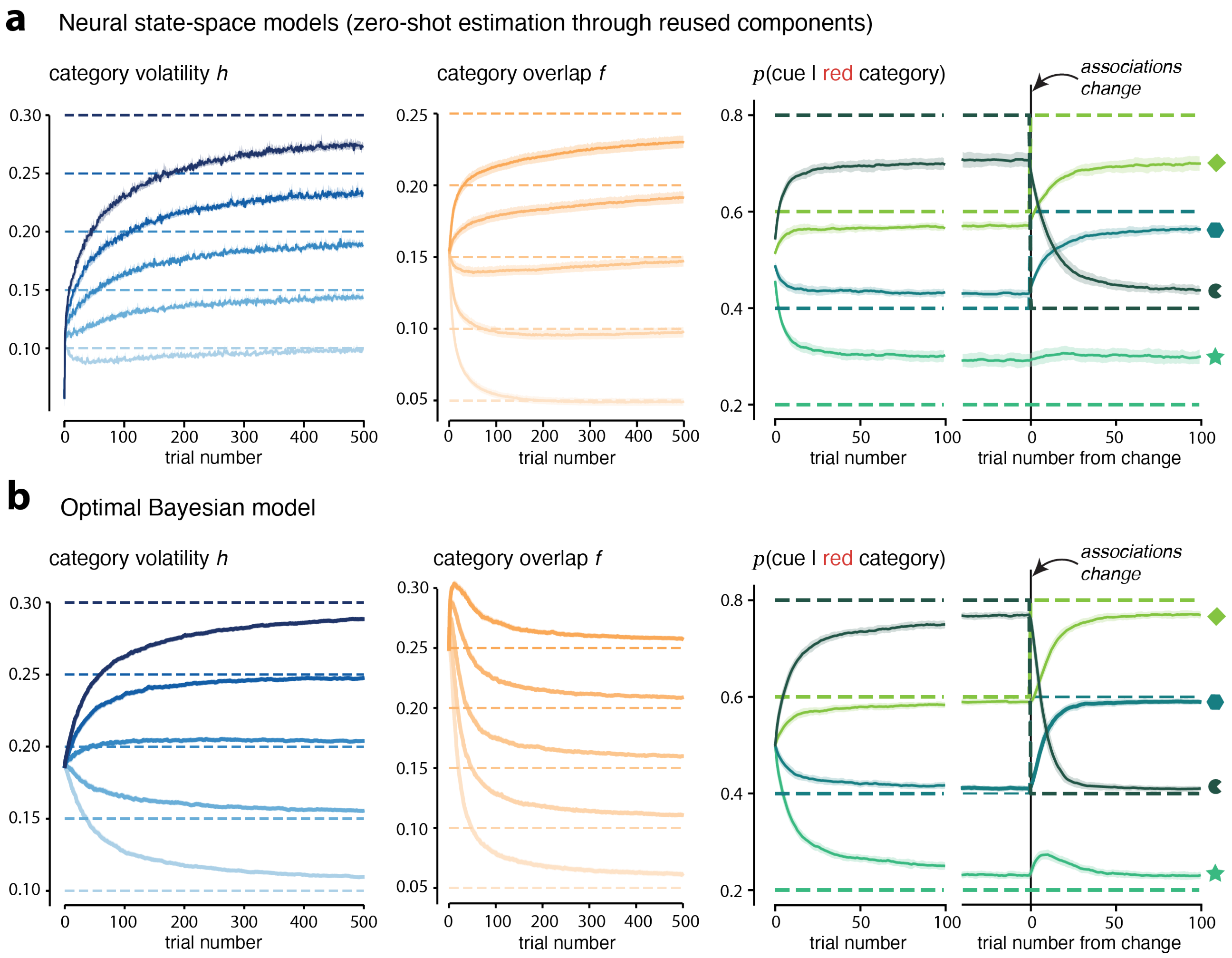


**Supplementary Figure 6 | Optimal Bayesian model in the history-dependent category prediction task** (**a**) Neural SSM estimates (reproduction of **Fig. 5c**). Left: Trial-by-trial estimates of the category volatility parameter $h$. Middle: Trial-by-trial estimates of the emission-overlap parameter $f$, reflecting uncertainty in the mapping from category to stimulus. Right: Time courses of estimated symbol–category associations, demonstrating flexible updating of the estimated associative probabilities following changes in associations. Importantly, these estimates are obtained in a zero-shot setting, with all components reused from previous tasks without any additional training. (**b**) Optimal Bayesian model. Same conventions as in (**a**). In contrast to neural SSMs, the optimal Bayesian observer is explicitly specified for this task and performs full task-specific inference over latent states and parameters. The optimal model shows qualitatively similar parameter dynamics, with convergence of volatility and emission-overlap estimates and flexible updating of symbol–category associations following changes in the underlying structure.

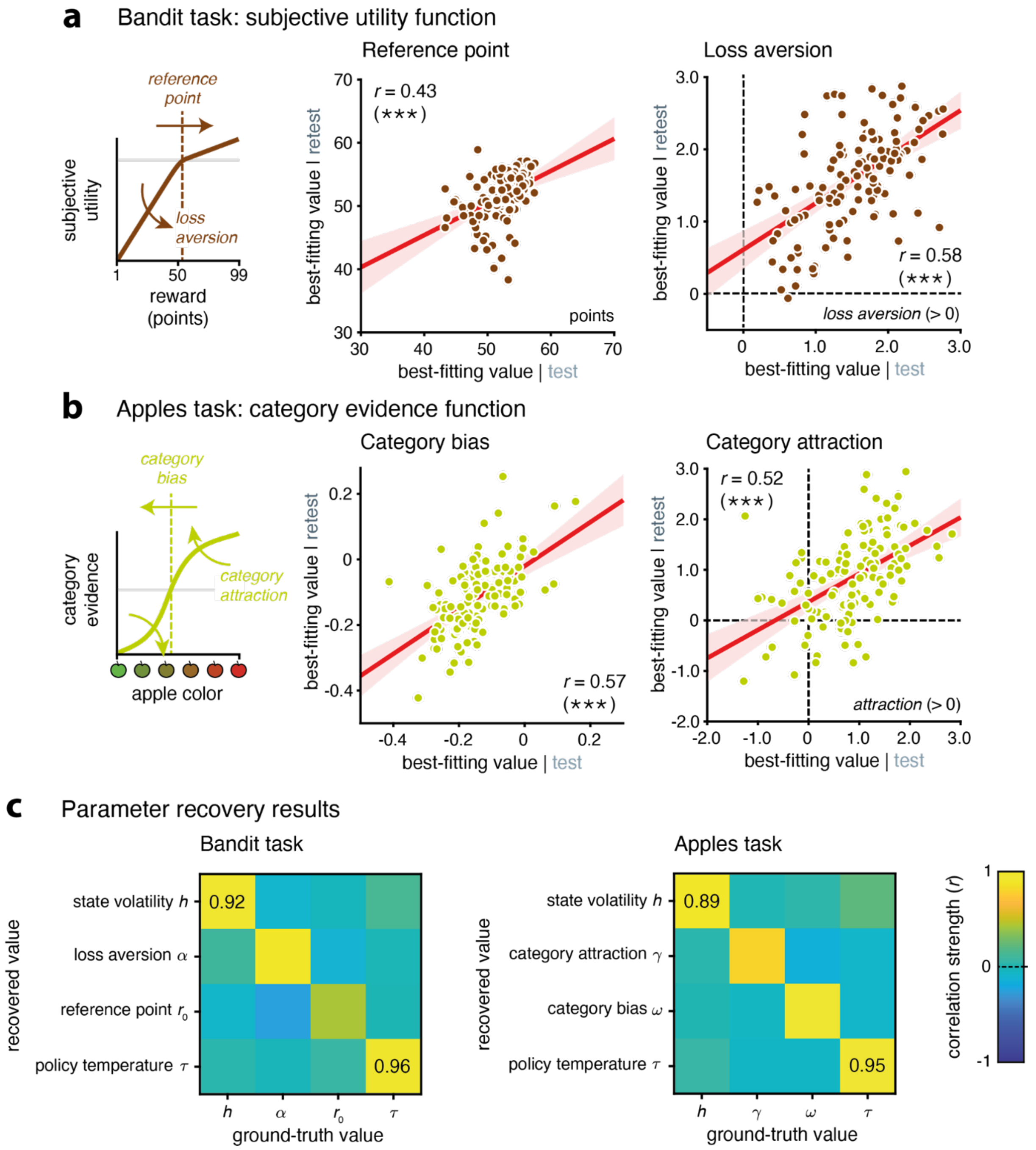


**Supplementary Figure 7 | Subjective distortions in the bandit and apples tasks** (**a**) Parameterization of the subjective utility function in the bandit task. Left: subjective utility function, controlled by a reference point and loss aversion (value > 0) or loss attraction (value < 0). Right: significant correlation of reference point (left) and loss aversion (right) across time in the bandit task. (**b**) Parameterization of the category evidence function in the apples task. Left: category evidence function, controlled by a category bias and category attraction (value > 0) or repulsion (value < 0). Right: significant correlation of category bias (left, toward red) and category attraction (right) across time in the apples task. Three stars indicate a significant effect at $p < 0.001$. (**c**) Parameter recovery for the hidden-state inference model. Parameter confusion matrices for the bandit (left) and apples (right) tasks, obtained by simulating behavior from the model using best-fitting parameters (estimated from human behavior), refitting the simulated data with the same model, and correlating recovered parameters with the original values. Recovery strengths (Pearson correlation coefficients) indicate robust parameter recovery, particularly for $h$ and $\tau$, which are the focus of our analyses.

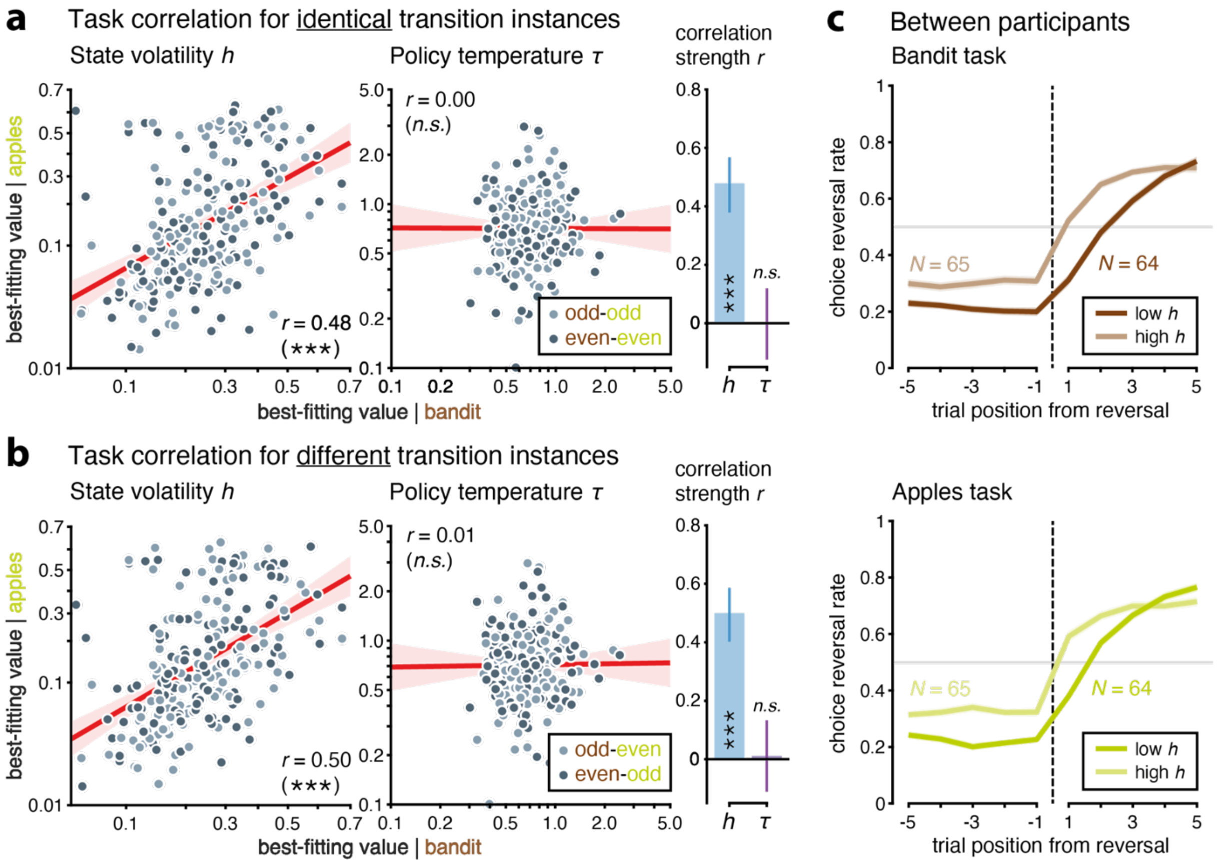


**Supplementary Figure 8 | Evidence of reuse across different transition instances in humans** (**a**) Correlation of parameters between tasks (x-axis: bandit task; y-axis: apples task) for identical transition instances, by comparing parameter fits to blocks with the same block number parity (odd-odd, even-even) across tasks. Left: strong correlation of state volatility $h$ (left) but zero correlation of policy temperature $\tau$ (right) between tasks. Right: correlation strength between tasks for h and τ. (**b**) Correlation of parameters between tasks for different transition instances, by comparing parameter fits to blocks with different block number parity (odd-even, even-odd) across tasks. Left: strong correlation of state volatility h (left) but zero correlation of policy temperature τ (right) between tasks. Right: correlation strength between tasks for $h$ and $\tau$. (**c**) Choice reversal curves for participants with low $h$ and participants with high $h$.

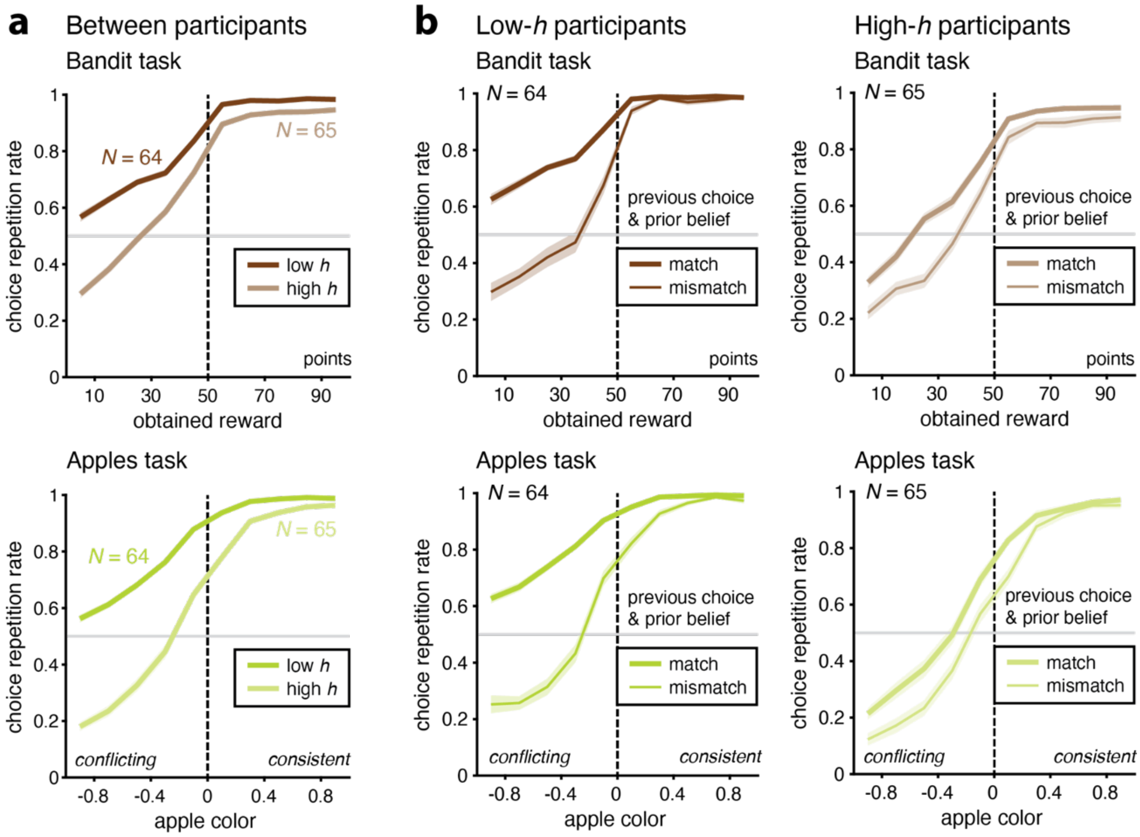


**Supplementary Figure 9 | Behavior signatures of individual differences in state volatility in humans** (**a**) Choice repetition rate as a function of obtained reward (bandit task, top) and apple color (positive: consistent with previous choice; negative: conflicting with previous choice, apples task, bottom) for participants with low (dark) and high (light) subjective state volatility $h$. Differences in $h$ are associated not only with differences in the overall choice repetition rate, but also with the sensitivity to the incoming evidence (obtained reward in the bandit task, apple color in the apples task). (**b**) Within-participant differences in choice repetition curve as a function of the match between the previous choice and the prior belief of the participant estimated using the inference model (thick: match; thin: mismatch), for participants with low (dark, left) and high (light, right) subjective state volatility $h$. In both tasks (top: bandit task; bottom: apples task), the match between the previous choice and the prior belief impacts choice repetition rate as a function of participants' subjective state volatility $h$.

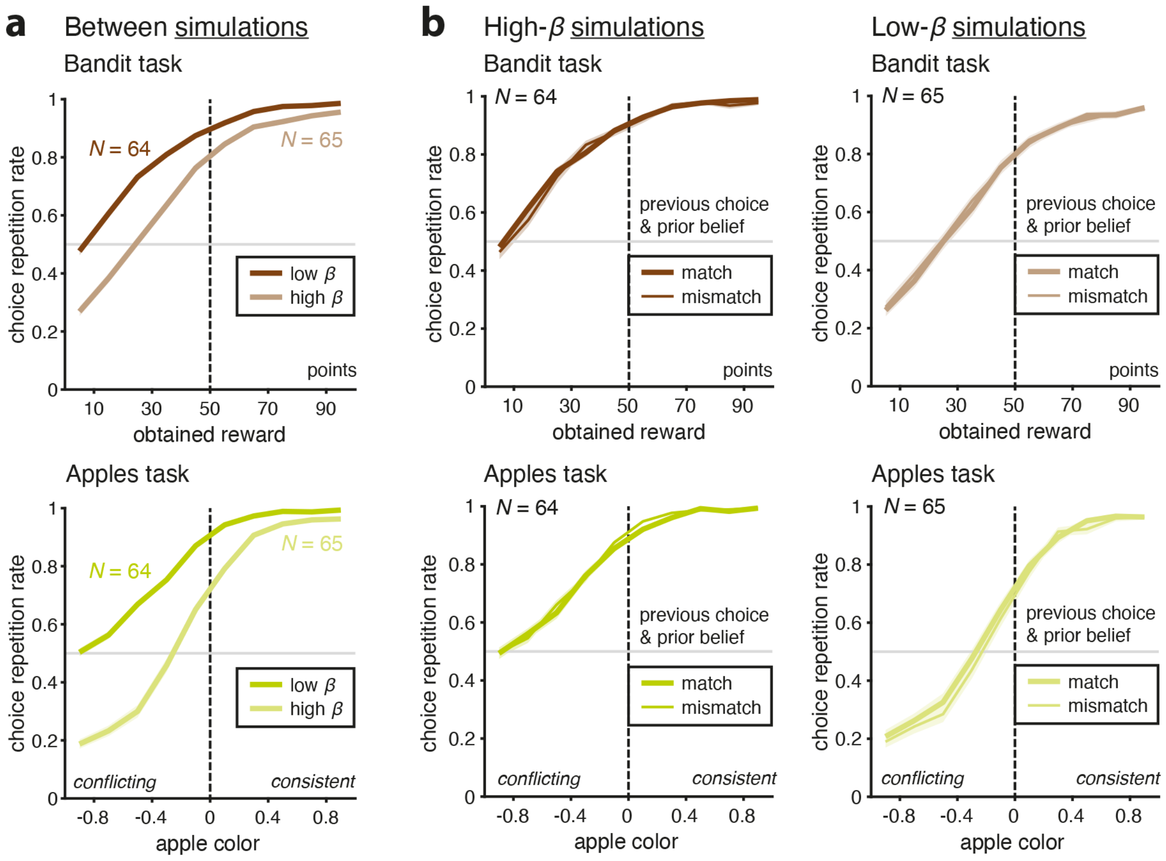


**Supplementary Figure 10 | Behavior signatures of individual differences in repetition bias in model simulations** (**a**) Choice repetition rate as a function of obtained reward (bandit task, top) and apple color (positive: consistent with previous choice; negative: conflicting with previous choice, apples task, bottom) for model simulations with low (dark) and high (light) repetition bias $\beta$ – but no transition component (i.e., $h$ = 0.5). Differences in $\beta$ across model simulations resemble differences in state volatility $h$ associated not only with differences in the overall choice repetition rate, but also with the sensitivity to the incoming evidence (obtained reward in the bandit task, apple color in the apples task). (**b**) Within-simulation differences in choice repetition curve as a function of the match between the previous choice and the prior belief estimated using the inference model with a subjective state volatility $h$ but no repetition bias $\beta$ (thick: match; thin: mismatch), for model simulations with high (dark, left) and low (light, right) repetition bias $\beta$. In both tasks (top: bandit task; bottom: apples task), the match between the previous choice and the prior belief does not impact choice repetition rate for model simulations with repetition bias but no transition component.

| | Computation time (mean ± SD, seconds) | |
|---|---|---|
| | **neural SSMs** | **Bayesian model** |
| Two-armed bandit | 0.64 ± 0.04 | 20.7 ± 1.3 |
| Stimulus prediction | 0.25 ± 0.02 | 140.4 ± 5.0 |
| Category prediction | 1.71 ± 0.05 | 606.9 ± 85.1 |
| History-dependent category prediction | 2.06 ± 0.09 | 731.6 ± 116.4 |

**Supplementary Table 1 | Computation time of neural SSMs and the optimal Bayesian model** Computation time (mean ± SD, in seconds) for neural state-space models (neural SSMs) and the Bayesian optimal model across four tasks. Each measurement was obtained by running 100 tasks in parallel, with each task consisting of 500 trials. For neural SSMs, computation times were averaged over 30 independently trained agents, each evaluated on a different instantiation of 100 parallel tasks. The optimal Bayesian model was evaluated on the same 30 task instantiations. Reported standard deviations reflect variability across these 30 runs.
